# Dynamic play between human N-α-acetyltransferase D and H4-mutant histones: Molecular dynamics study

**DOI:** 10.1101/2022.03.15.484466

**Authors:** Shravan B. Rathod, Kinshuk Raj Srivastava

**Affiliations:** Department of Chemistry, Smt. S. M. Panchal Science College, Talod, Gujarat, India; Division of Medicinal and Process Chemistry, CSIR-Central Drug Research Institute (CDRI), Lucknow, Uttar Pradesh, India

**Keywords:** Epigenetic modification, Histone acetylation, Oncohistone mutations, N-α-acetyltransferase D, Molecular docking, Molecular dynamics simulation

## Abstract

N-terminal acetyltransferases (NATs) are overexpressed in various cancers. Specifically in lung cancer, human N-α-acetyltransferase D (hNatD) is upregulated and prevents the histone H4 N-terminal serine phosphorylation, leading to the epithelial-to-mesenchymal transition (EMT) of cancer cells. hNatD facilitates histone H4 N-α-terminal serine acetylation and halts the CK2α-mediated serine phosphorylation. In the present study, we report the effects of four N-terminal mutant (S1C, R3C, G4D and G4S) histone H4 peptides on their bindings with hNatD by employing a molecular dynamics simulation. We also used graph theory-based analyses to understand residue correlation and communication in hNatD under the influence of WT and MT H4 peptides. Results show that S1C, R3C and G4S mutant peptides have significant stability at the catalytic site of hNatD. However, S1C, G4D and G4S peptides disrupt hNatD structure. Additionally, intramolecular hydrogen bond analysis reveals greater stability of hNatD in complex with R3C peptide. Further, intermolecular hydrogen bond analysis of acetyl-CoA with hNatD and its RMSD analysis in five complexes indicate that cofactor has greater stability in WT and R3C complexes. Our findings support previously reported experimental study on impacts of H4 mutations on its hNatD-mediated acetylation catalytic efficiency. The betweenness centrality (BC) analysis further gives insight into the hNatD residue communication dynamics that can be exploited to target hNatD using existed or novel drug candidates therapeutically.

**SECONDARY ABSTRACT:** Many N-terminal acetyltransferases (NATs) enzymes play important role in post-translational modification of histone tails. Research showed that these enzymes have been reported upregulated in many cancers. NatD is known to acetylate H4/H2A at the N-terminal. During lung cancer, this enzyme competes with the protein kinase CK2α and block the phosphorylation of H4 and, acetylates. Also, we observed that H4 has various mutations at the N-terminal and we considered only four mutations (S1C, R3C, G4D and G4S) to study the impacts of these mutations on H4 binding with NatD using MD simulation. Our results show that R3C stabilizes the NatD whereas remaining mutations destabilize the NatD. Thus, mutations have significant impacts on NatD structure. Our finding supports previous analysis also.

**SIGNIFICANCE:** Our main objective in this study was to understand the structural and dynamics of hNatD under the influence of WT and MT H4 histones bindings. Previous experimental study reported that mutations on H4 N-terminus reduce the catalytic efficiency of N-Terminal acetylation. But here, we performed molecular-level study thus, we can understand how these mutations (S1C, R3C, G4D and G4S) cause significant depletion in catalytic efficiency of hNatD. Another, interesting observation is that enzymatic activity of hNatD is altered due to the considerably large deviation of acetyl-CoA from its original position (G4D). Further, simulation and correlation data suggest which regions of the hNatD are highly flexible and rigid and, which domains or residues have the correlation and anticorrelation. As hNatD is overexpressed in lung cancer, it is an important drug target for the cancer hence, our study provides structural information to target hNatD.

## INTRODUCTION

Oncogenesis is complex and multistep process involves mutations in epigenetic enzymes. Current experimental evidences indicate that histones are found to be mutated in numerous cancers. For instance, histone H3 was reported mutated in rare paediatric gliomas and sarcomas (1). Additionally, in large B-cell lymphomas, histone H1 was found mutated (2). Nacev et al. reported cancer associated histone mutations recently (3).

Epigenetic histone tail modifications portray vital role in gene expression at the transcription site for the protein synthesis. While modification at the nucleosome region facilitates the chromatin condensation and regulates various nuclear events (4). Among the phosphorylation, methylation, acetylation, deacetylation, ubiquitylation, and sumoylation post-translational modifications (PTMs), acetylation and methylation have been widely studied (5). N-α-terminal acetylation is most frequently occurring covalent modification in approximately 90% soluble proteins in humans and around 70% in yeast (6-9). This covalent modification is involved in cellular metabolism, apoptosis, protein translocation, membrane attachment, protein-protein interactions, protein regulation and degradation (9–12).

N-α-acetyltransferases (NATs) catalyse N-terminal acetylation of alpha-amino group of protein by using acetyl-coenzyme A (Ac-CoA). There have been total six NATs reported in humans according to their specific subunits and substrates (9). hNatD is also known as Patt1 (protein acetyltransferase) or Nat4 exclusively acetylates the histone H2A and H4 compared to other NATs those target numerous other substrates (13–15). Ju et al. have recently reported that hNatD is overexpressed in lung cancer and it elevates the cancer progression by blocking the protein kinase 2-alpha (CK2α)-mediated histone H4 serine phosphorylation. hNatD acetylates the histone H4 serine at alpha-amino position competitively and increases the rate of epithelial-to-mesenchymal transition (EMT) of cells in lung cancer (16). The hNatD acetylates the N-terminus residues of H4 and H2A by using acetyl-CoA cofactor located inside the catalytic tunnel in hNatD. Fig. 1 illustrates the acetylation of H4/H2A by hNatD. Recently, how the activity and catalytic efficiency of hNatD are altered under the influence of WT and MTs pentapetides binding to hNatD has been reported (17). However, the molecular level understanding of H4 (WT & MTs) binding dynamics still lacks.

**FIGURE 1.**
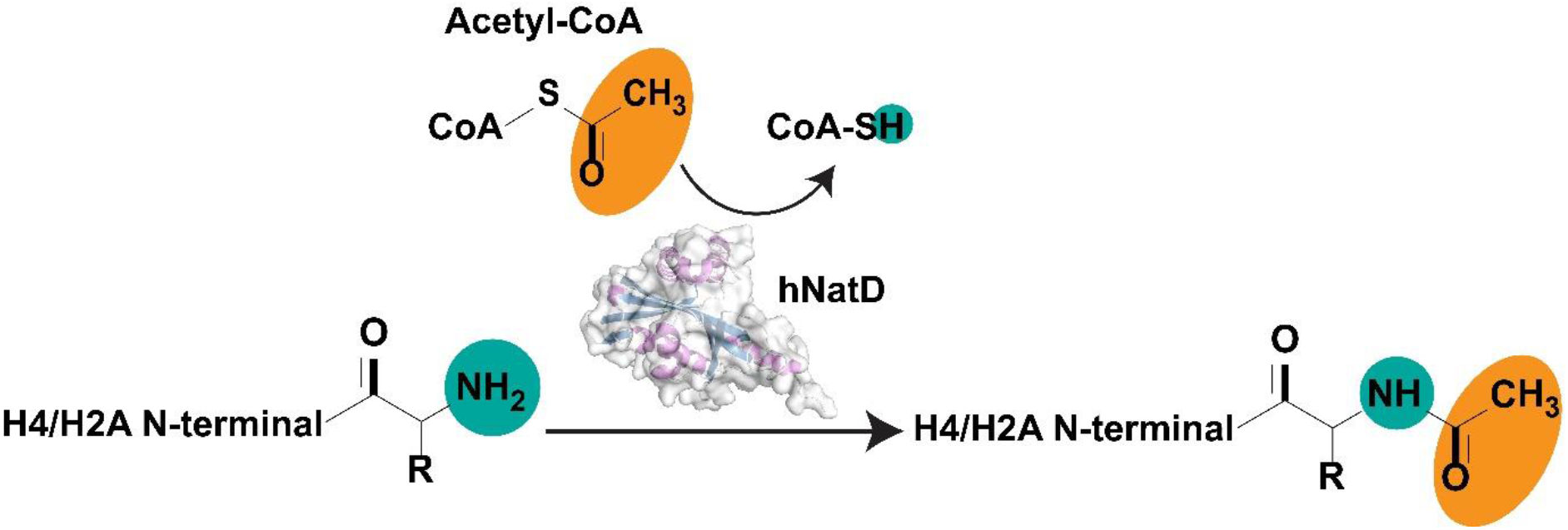
N-terminal acetylation of H4/H2A histone by human N-α-acetyltransferase D (hNatD) enzyme using acetyl-CoA cofactor.

In the present study, our aim was to investigate the impacts of four mutations at the histone H4 peptide on its binding at the hNatD acetylation site and, to probe the structural and dynamic characteristics of hNatD to target it therapeutically in the future. We used molecular dynamics simulation (MDS) approach to probe the binding events of hNatD and histone H4 (WT and MTs) tails. Additionally, residue interaction network (RIN) based analyses were carried out to further understand hNatD structural dynamics.

## MATERIALS AND METHODS

### PROTEIN PREPARATIONS

The crystal structure (PDB: 4U9W) (13) of hNatD (a.a.: 1-207) with histone H4 peptide and acetyl-CoA was retrieved from the protein data bank (PDB). SWISS-MODEL (18) available at http://swissmodel.expasy.org was employed to add missing residues in the β6-β7 loop and N-terminal α0-helix in hNatD. Additionally, the missing acetyl group in acetyl-CoA was added using Builder tool in PyMOL 2.4.1 (19).

### PROTEIN-PEPTIDE DOCKING

In the crystal structure, histone H4 peptide had five residues (SGRGK). To fully investigate the impacts of four mutations (S1C, R3C, G4D, and G4S) of histone H4 peptide on its binding with hNatD, we wanted to add another five residues to the H4 peptide. Thus, the HPEPDOCK protein-peptide docking web server (20) available at http://huanglab.phys.hust.edu.cn/hpepdock/ was employed in which modelled hNatD was uploaded as a receptor and for ligand (H4 peptide), 10 aa short sequence (SGRGKGGKGL) of H4 peptide was provided in the ligand box. In the case of sequence query, HPEPDOCK utilizes MODPEP program (21) to generate ensembles of peptides and further these ensembles of peptides are docked with receptor using MDock program suite (22), in which rigidity is applied to the receptor and flexibility is given to peptide during the docking. To determine the 3D interactions of H4 peptide with hNatD, Discovery Studio v.20.1 (23) was used.

### MOLECULAR DYNAMICS SIMULATION

Molecular dynamics simulation was employed to scrutinize binding between the histone H4 (WT & MTs) and hNatD. Four mutations in histone H4 peptide were inserted using Mutagenesis tool in PyMOL. These four mutants and wildtype complexes were further subjected to the 100 ns molecular dynamics (MD) simulation. Molecular dynamics simulation was carried out using GROMACS 2020.1 version (24). The updated version of CHARMM36 force filed (charmm36-feb2021) (25,26) and TIP3P water model (27) were employed. To build the acetyl-CoA topology, CHARMM general force field (CGenFF) (28, 29) available at (https://cgenff.umaryland.edu/) was utilized. For the simulations, these four complexes were placed at the center inside the cubic box with a 10 Å from the box edges. Further, positive (Na^+^) and negative (Cl^−^) ions were added to neutralize the solvation system. Initially, the system was energy minimized using the steepest descent algorithm (30) to reduce the steric clashes between the atoms. For the optimization of long-range electrostatic forces, Verlet cut-off (31) and particle mesh Ewald (PME) schemes (32) were employed. During the energy minimization protocol, nsteps was set to 50,000 and energy cut off was kept 10 kJ/mol. Energy minimization was followed by two subsequent equilibrations (NVT and NPT) at 300 K temperature for 100 ps (50,000 steps) with a 2 fs time step gap. Then, MD simulations was carried out for the 100 ns using leap-frog integrator (33), Verlet cut-off scheme, and PME at 300 K (modified Berendsen thermostat (34)) and 1 bar pressure (Parrinello-Rahman method (35)). LINCS (linear constraint solver) algorithm (36) was utilized to constrain bonds and angles. CHARMM36 force filed and python script were downloaded from the MacKerell lab website (http://mackerell.umaryland.edu/charmm_ff.shtml#gromacs). Finally, the MD trajectories were analyzed for the root mean square deviation (RMSD), solvent accessible surface area (SASA), radius of gyration (Rg), distances, root mean square fluctuation (RMSF) and, inter and intramolecular hydrogen bonds in the complexes.

### PROTEIN-PEPTIDE INTERACTIONS ANALYSIS

To investigate interactions between histone H4 and hNatD, ten frames from the 100 ns each simulation were extracted at every 10 ns from 10 ns to 100 ns. The web-based tool, PDBePISA (https://www.ebi.ac.uk/pdbe/pisa/) was employed to calculate numbers and types of interactions in 10 frames of each four complexes; hNatD-H4_WT, hNatD-H4_S1C, hNatD-H4_R3C, hNatD-H4_G4D and hNatD-H4_G4S. We considered dominant interactions in 10 frames of each complex and they are illustrated. PDBePISA calculates interface area and number of interacting residues between the proteins and solvation energy along with interactions. To calculate van der Waals, hydrogen bond, electrostatic and stabilization energies of WT and MT complexes, PIMA web server (37) available at http://caps.ncbs.res.in/pima/ was also employed.

### LINEAR MUTUAL INFORMATION

To probe the dynamical behaviour and fluctuations of proteins, dynamical cross-correlation (DCC) and linear mutual information (LMI) are used widely (38-41). Considering the limitation of DCC that it does not calculate correlation of simultaneously moving atoms in perpendicular directions (42), we utilized normalized LMI (nLMI). To perform the nLMI on complexes, we used recently developed python 3-based tool, *correlationplus* (42). To calculate nLMI, we provided 100 ns MD trajectory files (XTC format) as an input to the program. From the 100 ns trajectories, it used 10,000 frames from each trajectory to calculate nLMI. Additionally, we calculated residue betweenness to find significant correlated residues. Commands to run *correlationplus* is given in Table S1.

### ELECTROSTATIC POTENTIAL (ESP)

The eF-surf web server available at https://pdbj.org/eF-surf/top.do was used to calculate electrostatic potential surface of hNatD. This server requires only protein structure file in pdb format as an input file. Avogadro (43) was used to generate ESP surface of the acetyl-CoA while PyMOL APBS Electrostatics plugin was used to generate ESP of histone H4 WT and MT peptides.

## RESULTS AND DISCUSSION

### STRUCTURAL FEATURES OF hNatD

The structural features of hNatD are resemble to the previously reported other NATs (N-acetyltransferases) and it contains α/β mixed GCN5-like fold (Fig. 2) (44–46). From the comparison of hNatD with NatA and NatE, it was observed that hNatD has some specific additional structural features which are loops formed by α1-α2 and β6-β7 and, unique N-terminal helix (α0). The hNatD sequence alignments among various organisms revealed that a specific region, 55-216 a.a. (~150 residues) is highly conserved in hNatDs. For the substrate recognition by NatA and NatE, they have β6-β7 hairpin loop region. However, there is significant structural difference observed at the catalytic site of hNatD. In hNatD, β6-β7 loop is located slightly away from the substrate binding site (13).

**FIGURE 2.**
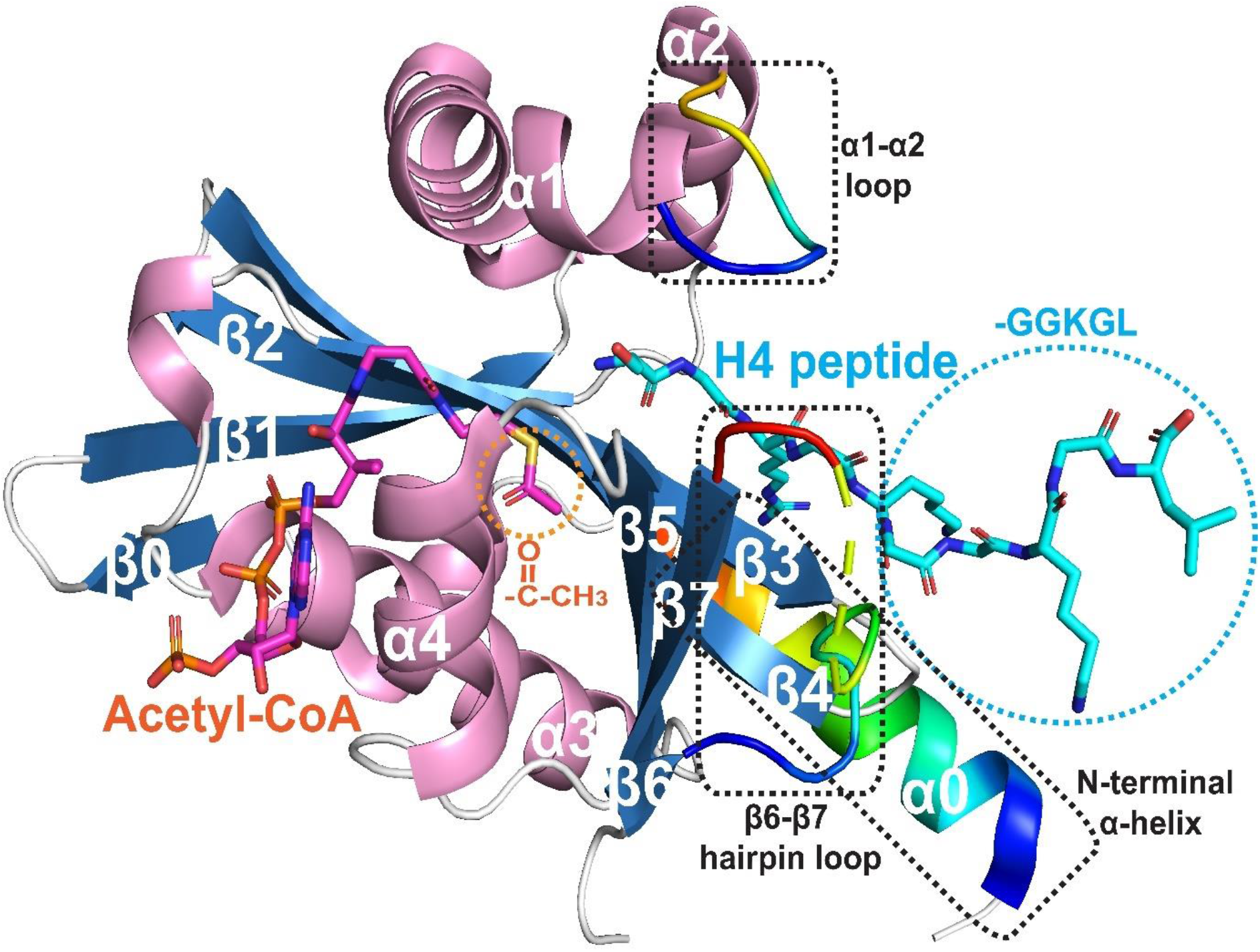
Structure of hNatD with acetyl-CoA and histone H4-tail (PDB: 4U9W). In this crystal structure, acetyl group was missing that was added and another five amino acids were added to the H4 peptide. Circular dashed rings indicate the modification sites. Three important structural features of hNatD, α1-α2 loop, β6-β7 loop and N-terminal α-helix are represented in rainbow color and shown in rectangular dashed boxes.

In addition to this, α1-α2 loop further protects catalytic region. Another interesting feature is helix-loop-strand forming N-termini region (24–51 a.a.). This region stabilizes the whole hNatD by wrapping around the hydrophobic patch of substrate recognition site. Five main residues, P38, L39, F42, F45 and Y48 of N-terminal interact with the L53, I57, C59, W107, L109, F124, F126, L151, F154, I158, L161 and M162 residues of catalytic site. These observations indicate that hNatD has specific structural features to recognize histone H4 or H2A.

The electrostatic potential (ESP) surface analysis is crucial to understand molecular mechanisms such as intermolecular association of protein and small ligands, actions of drug candidate, enzyme catalysis and protein-DNA interactions complementarity (47). In this study, we also calculated electrostatic potential surface of the hNatD. It can be seen from Fig. 2, left-hand side on hNatD has positively charged residues (R134, R135, K136 and K140) containing cleft which harbours acetyl-CoA (Fig. 3A). As acetyl-CoA has negatively charged phosphate backbone (Fig. 3C), it perfectly positioned itself into this cleft. In contrast to that on the right-hand side on hNatD, histone H4 binding site has negatively charged residues (E76, E83, E86, E87, D90, D114, E116, E126 and E194) which recognises positively charged histone H4 peptide (Fig. 3B). Histone H4 peptide (SGRGKGGKGL) has arginine and lysine basic residues which make this surface positively charged (Fig. 3F). hNatD exhibits long tunnel to harbour acetyl-CoA and histone H4 peptides (Fig. 3D-E).

**FIGURE 3.**
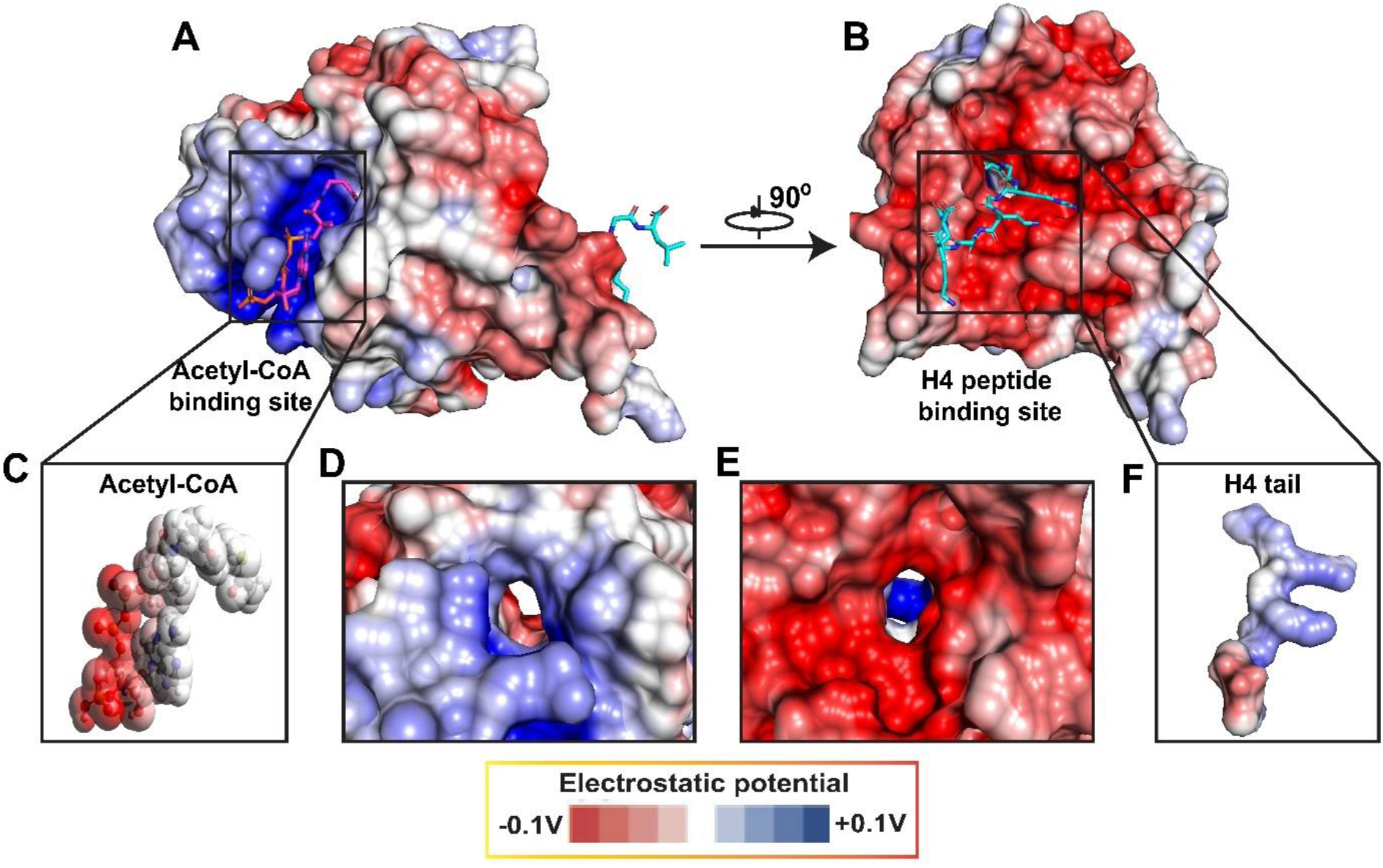
Electrostatic potential (ESP) surface representations. (A) Positively charged binding site of acetyl-CoA. (B) Negatively charged binding site of histone H4 peptide. (C) ESP of acetyl-CoA (D) Catalytic tunnel from acetyl-CoA binding site to H4 peptide binding site. (E) Catalytic tunnel from H4 peptide binding site.to acetyl-CoA binding site. (F) ESP of histone H4 peptide.

### PROTEIN-PEPTIDE DOCKING

Molecular docking is used widely to investigate the bindings of protein with protein, peptide, small ligand, DNA and RNA in computer-aided drug design (CADD) and other computational analyses to complement experimental findings (48–54). In this study, to probe the impacts of four mutations (S1C, R3C, G4D, and G4S) in histone H4 peptide (10 a.a.), we first docked histone H4 peptide with hNatD using HPEPDOCK protein-peptide web server (20). From the top 100 predictions, best model was selected based on docking score (−172.891 kcal/mol). This docked model was structurally aligned with crystal structure (PDB: 4U9W) in PyMOL and RMSD was found 0.022 nm (Fig. S1). Further, we obtained protein-peptide 3D interaction profile using Discovery Studio v.20.1 tool. Fig. 4 illustrates best docking pose (Fig. 4A) and 3D interactions between the hNatD and histone H4 peptide (Fig. 4B).

**FIGURE 4.**
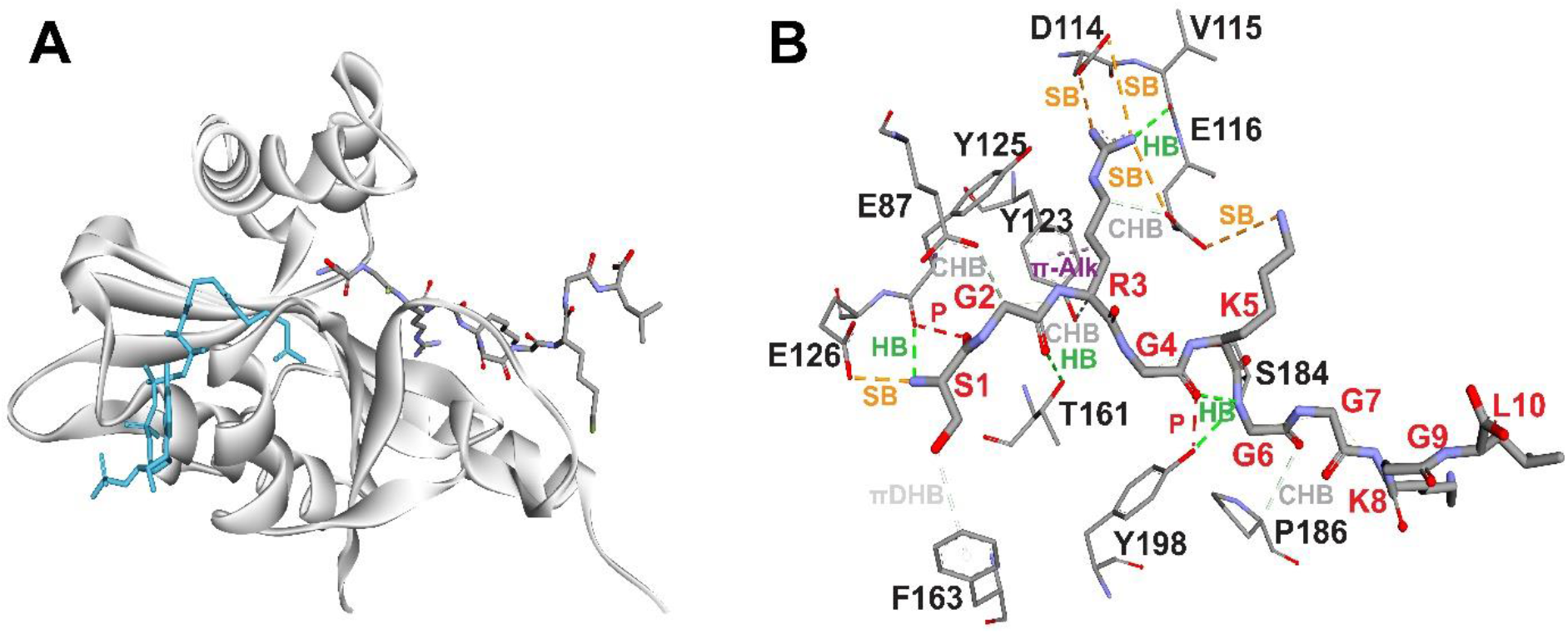
(A) H4 peptide docked structure. The acetyl-CoA and H4 peptide are shown in stick representation and have cyan and grey colors respectively. (B) 3D-interactions between hNatD and H4 peptide. H4 peptide amino acids’ labels are colored in red and hNatD in black. HB: hydrogen bond, SB: salt bridge, CHB: carbon hydrogen bond, π-Alk: pi-alkyl bond, πDHB: pi-donor hydrogen bond and P: polar.

It can be seen from the Fig. 3B that histone H4 peptide is surrounded with mainly acidic and aromatic residues. Histone tail forms salt bridges with the acidic residues and hydrogen bonds with other residues of the hNatD. And, two aromatic residues (Y123 and F163) from the hNatD interact through donating pi electrons to the peptide residues. Additionally, polar interactions (carbon-hydrogen bond) can be observed between hNatD and histone H4 peptide.

### IMPACTS OF HISTONE H4 PEPTIDE BINDING ON THE DYNAMICS OF hNatD

To target hNatD therapeutically, histone H4 interacting residues and their communication are significant. Thus, we analysed the impacts of WT and MT histone H4 decapeptides bindings on dynamic behaviour of hNatD through employing MD simulation and nLMI analyses. Ligand binding to protein induces structural alternations in proteins (55–57). To understand which regions of the hNatD fluctuate largely, we aligned MD trajectory frames from 0 ns to 100 ns at every 10 ns for each case (Fig. 5). The average RMSDs of hNatD after aligning frames in each complex were observed 0.091 nm, 0.091 nm, 0.087 nm, 0.118 nm and 0.085 nm for hNatD-H4_WT, hNatD-H4_S1C, hNatD-H4_R3C, hNatD-H4_G4D and hNatD-H4_G4S respectively. Among these five complexes, hNatD-H4_R3C (Fig. 5C) and hNatD-H4_G4S (Fig. 5E) showed considerably less deviation in comparison with other complexes (Fig. 5A-B, D). Radar plots illustrate the H4 head and tail dynamics in all five complexes (Fig. 5F-G). In radar plots, distances of Cα-atom of first residue in every frame (10 ns-100 ns at every 10 ns) with respect to Cα-atom of first residue in 0 ns frame were calculated and plotted. It can be observed from the Fig. 5F that heads of WT and G4D histones showed greater deviation (2 nm-2.5 nm) from their original positions at 0 ns frame in comparison with S1C, R3C and G4S MTs histones which showed at around 0.5 nm-1.0 nm. However, in case of H4 histones, distances of Cα-atom of last residue (L10) in every frame (10 ns-100 ns at every 10 ns) with respect to Cα-atom of last residue in 0 ns frame were calculated and plotted. However, in this case, except R3C, every histone showed large fluctuations and deviations were around 1.5 nm-2.5 nm (Fig. 5G).

**FIGURE 5.**
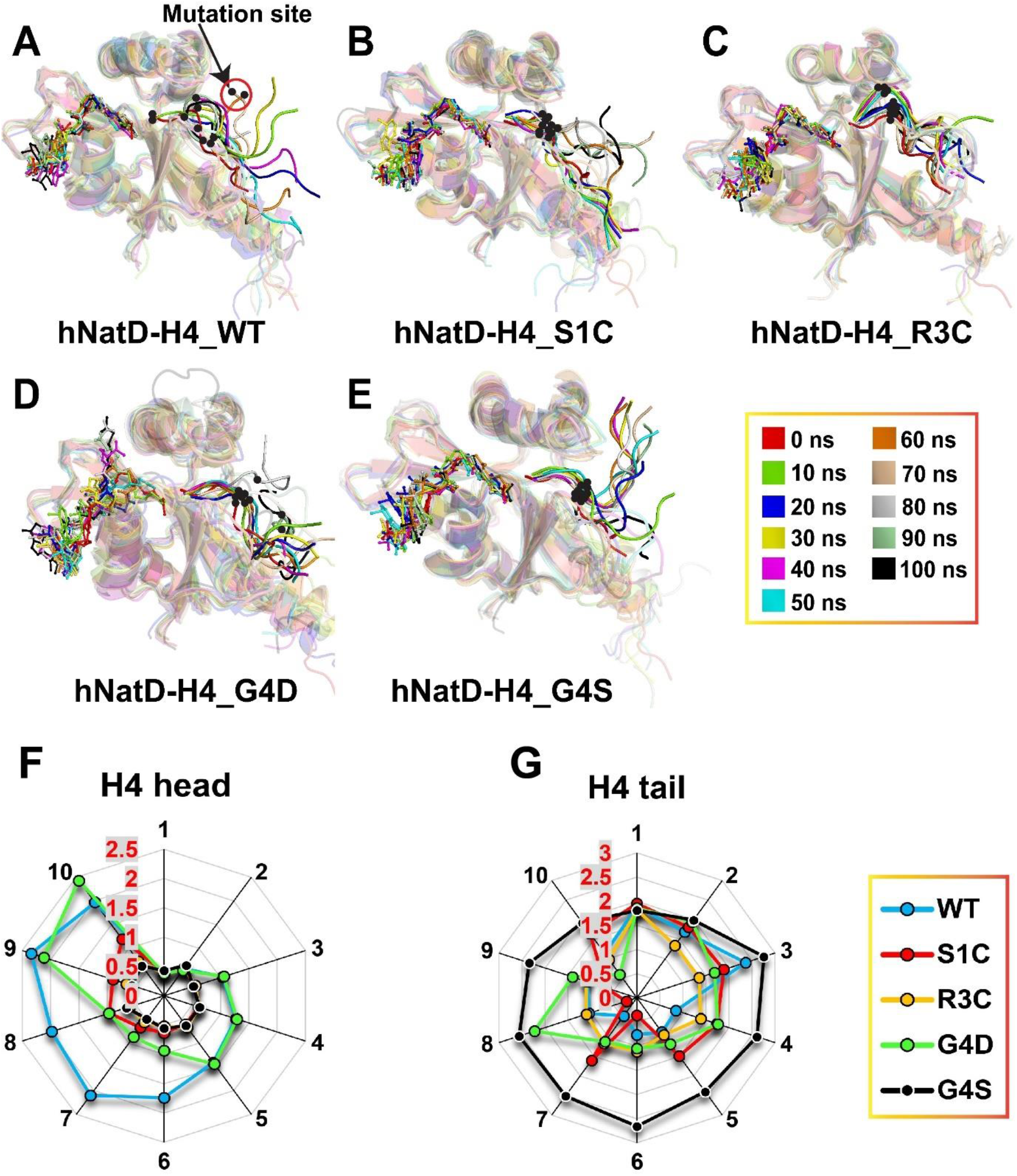
Structural alignments of 100 ns MD simulation frames at every 10 ns from 0 ns to 100 ns. (A) hNatD-H4_WT complex. (B) hNatD-H4_S1C complex. (C) hNatD-H4_R3C complex. (D) hNatD-H4_G4D complex. (E) hNatD-H4_G4S complex. (F) Radar plot of histone H4 (WT & MTs) heads’ movements. (G) Radar plot of histone H4 (WT & MTs) tails’ movements. In radar plots, labels on circular path indicate the number of frame (1-10) whereas vertical values (red color) are distance ranges (0-3 nm) between H4 heads or tails frames. Black dots designate the mutation position in H4 peptide.

To understand the histone H4 peptide behaviour inside the catalytic site of hNatD, we also calculated the RMSDs of WT and MTs histone H4 peptides (Fig. 6A). Mutant histone H4_G4S showed larger deviation (RMSD: 1.21 nm) from the initial position which is responsible to the higher flexibility of its tail. The average RMSDs of H4_WT, H4_S1C, H4_R3C and H4_G4D histone H4 peptides were observed 0.96 nm, 0.97 nm, 0.88 nm and 0.94 nm respectively. Thus, except H4_R3C, all the WT and MTs H4 peptides have greater dynamicity inside the hNatD cavity. We also determined dynamics of acetyl-CoA inside the hNatD cavity under the influence of histone H4 WT and MTs tails bindings to the hNatD. Fig. 6B shows the RMSD of acetyl-CoA in hNatD of all five complexes. The acetyl-CoA showed greater fluctuations (0.54 nm) inside the binding pocket in hNatD of hNatD-H4_G4D as compared to other complexes. The average RMSDs of acetyl-CoA in hNatD-H4_WT, hNatD-H4_S1C, hNatD-H4_R3C and hNatD-H4_G4S were found 0.24 nm, 0.28 nm, 0.24 nm and 0.30 nm respectively. To find the cause of this trend, we looked at the electrostatic surface of the WT and MTs H4 histones (Fig. S2) and hNatD at the binding pocket (Fig. 3B). The histone H4 binding cavity has the dominant glutamic acidic residues which make pocket negatively charged surface. Except the H4_G4D, all histones have positively charged surface (Fig. S2). In case of H4_G4D, glycine smaller amino acid is substituted with acidic aspartic acid residue which carries negatively charged carboxylic group (Fig. S2(D)). Thus, due to the repulsion between the H4_G4D and hNatD catalytic cleft, this mutation significantly disrupts the residue network in hNatD_H4_G4D and also the acetyl-CoA fluctuates largely (Fig. 5(D)).

**FIGURE 6.**
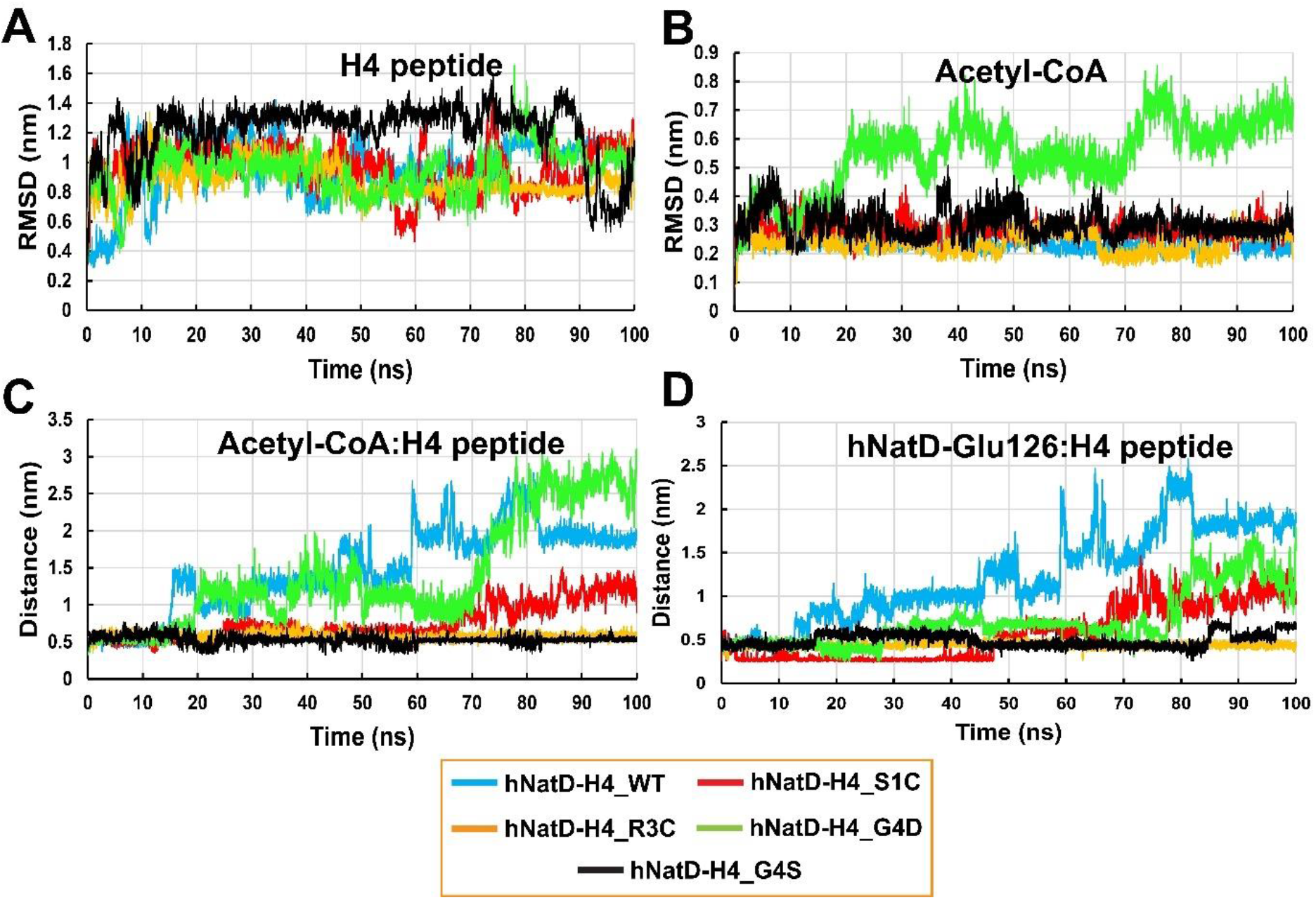
(A) RMSD of histone H4 peptides (WT & MTs). (B) RMSD of acetyl-CoA. (C) Distance between acetyl-CoA sulfur (S) atom and nitrogen (N) atom of amino group in residue 1 (Serine/Cysteine) of histone H4 peptides (WT & MTs). (D) Distance between hNatD E126 carboxylic oxygen (O) atom and nitrogen (N) atom of amino group in residue 1 (Serine/Cysteine) of histone H4 peptides (WT & MTs).

The hNatD acetylates the histone H4 and H2A N-terminals using acetyl-CoA cofactor (13). For the acetylation, the distance between the amino group of residues in histones and sulfur atom of acetyl-CoA must be in close proximity. Thus, we calculated distances between amino group nitrogen of S1/C1 and sulfur atom of cofactor in WT and MTs complexes (Fig. 6C). In WT and G4D, the position of the first residue deviated greatly from its original position thus, distances were observed significantly large (~0.5-2.5 nm) during the 100 ns simulation. However, first residue in S1C and G4S was observed highly stable throughout the simulation and average distance was noted 0.5 nm. In R3C, distance was observed approximately 0.5 nm until 65 ns and then, it was increased and reaches to the around 1.5 nm at the end of the simulation. Additionally, E126 residue in hNatD was confirmed to be important to catalyse the acetyltransferase reaction not by directly acting as deprotonating agent but by increasing the nucleophilicity of S1/C1 amino group through inducing dipole moment from the carboxylic group. Also, it is believed that this residue orients the S1/C1 amino group towards the acetyl-CoA to facilitate reaction and it is considered essential for the catalysis (13). Thus, we calculated the distance between the carboxylic oxygen (-CO(OH)) of E126 and amino group nitrogen (N) atom of S1/C1 histone tail (Fig. 6D). It can be observed that except WT, in all other complexes, until around 65 ns, distance was remained approximately 0.5 nm. And, more importantly, it was remained stable during the entire simulations of hNatD-H4_R3C and hNatD-H4_G4S complexes. Hence, this data suggests that E126 is vital residue for the catalysis and also validate the previous finding.

After performing 100 ns simulations, the dynamic behaviour of hNatD was determined using various post-MD trajectory analyses. The Root mean square deviation (RMSD) of hNatD in five complexes (hNatD-H4_WT, hNatD-H4_S1C, hNatD-H4_R3C, hNatD-H4_G4D and hNatD-H4_G4S) was calculated using backbone atoms to understand the conformational changes in hNatD under the influence of histone H4 peptide (WT & MTs) binding. The RMSD can be calculated using following equation;

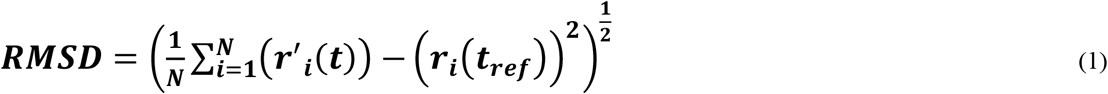

Where, *N* indicates the total number of particles (atoms), *r_i_* is the position of atom *i* at reference time *t*_*ref*_, *r*′ is the position of the selected atom at time *t* after aligning with reference frame.

Fig. 7A shows the RMSD plot of hNatD in studied five complexes. The average RMSD of hNatD in hNatD-H4_WT, hNatD-H4_S1C, hNatD-H4_R3C, hNatD-H4_G4D and hNatD-H4_G4S was observed 0.025 nm, 0.025 nm, 0.24 nm, 0.28 nm and 0.27 nm respectively. Thus, data indicates that hNatD has more structural deviations and fluctuations in hNatD-H4_G4D and hNatD-H4_G4S as compared to others. It can be seen from the RMSD plots also in Fig. 7A. The hNatD in hNatD-H4_R3C has the highest stability (RMSD: 0.24 nm) and throughout the 100 ns simulation, RMSD oscillates between 0.20 nm and 0.25 nm. However, in case of hNatD-H4_G4D and hNatD-H4_G4S, there were significant changes noted during the simulation and RMSDs were fluctuated between 0.15 nm and 0.43 nm. Fig. S3-S7 shows hNatD regions which are significantly dynamic in nature. In all the complexes, mainly four regions, N-terminal α0-helix (~1-20 a.a.), α1-α2 loop (~68-81 a.a.), α4-helix (~166-176 a.a.) and β6-β7 loop (~180-198 a.a.) exhibit strong deviation except in case of hNatD-H4_R3C, in which only β6-β7 loop showed deviation. Additionally, α2-helix (~81-90 a.a.) in hNatD-H4_G4S showed deviation addition to remaining four regions compared to others.

**FIGURE 7.**
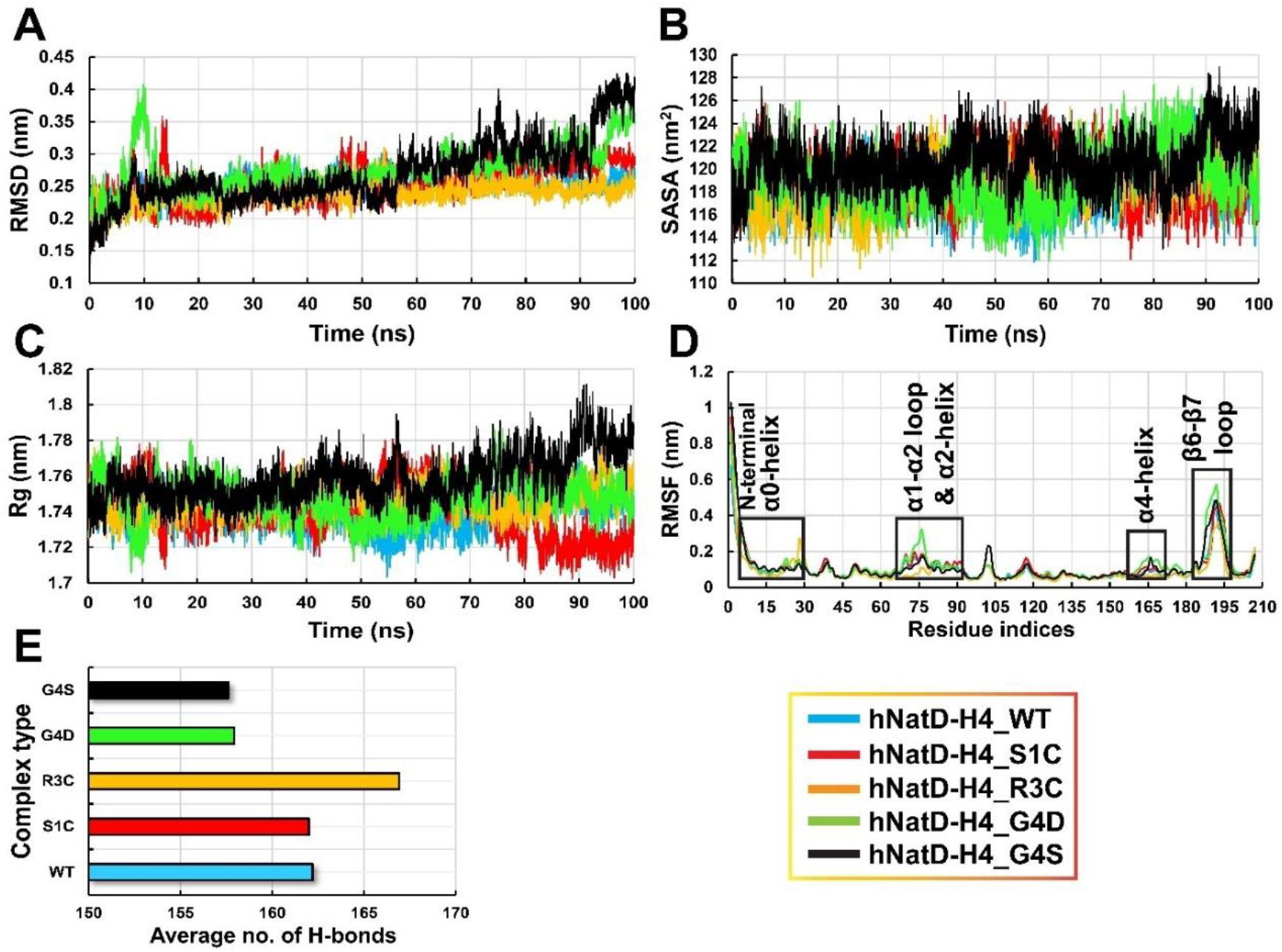
MD trajectory post analyses of five complexes (WT & MTs). (A) RMSDs (nm) plots of hNatD. (B) SASA (nm2) plots of hNatD. (C) Rg (nm) plots of hNatD. (D) RMSF (nm) plots of hNatD. (E) Average intramolecular hydrogen bonds in hNatD.

From the RMSD plots in Fig. 7A, it can be noted that there is a peak (0.40 nm) at 10 ns and then RMSD was decreased to normal (0.25 nm) in hNatD-H4_G4D. Afterwards it showed gradual increment and reached to the ~0.35 nm at the end of simulation. However, hNatD in hNatD-H4_G4S showed four significant changes in RMSD during 0-25 ns, 26-55 ns, 56-90 ns and 91-100 ns times.

Moreover, solvent accessible surface area (SASA) and radius of gyration (Rg) were calculated to investigate the hNatD folding and compactness. The SASA and Rg give similar trend in the plots. Both indicate the opening or closing of hNatD during the simulation. The SASA is the total surface area which is in contact with solvent (water) whereas, Rg is the average distance of each atom from the centre of gravity in protein. The Rg is calculated using below equation;

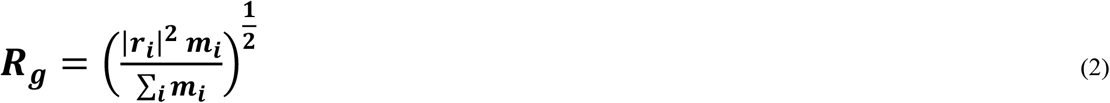

Where, *r_i_* and *m_i_* are the position and mass of the atom *i* respectively in protein. The average SASA values of hNatD in hNatD-H4_WT, hNatD-H4_S1C, hNatD-H4_R3C, hNatD-H4_G4D and hNatD-H4_G4S were found 118.3 nm^2^, 119.2 nm^2^, 118.9 nm^2^, 119.2 nm^2^ and 120.6 nm^2^ respectively. The higher value of SASA or Rg indicates the reduction in protein compactness and lower value suggests gain in the compactness. It can be observed from the SASA (Fig. 7B) and Rg (Fig. 7C) plots that the folding and compactness of hNatD in hNatD-H4_WT and hNatD-H4_G4D are following similar trend. Initially, the SASA values gradually decline from around 121 nm^2^ to 116 nm^2^ until ~55 ns. After that both the values begin to rise and reach to approximately 124 nm^2^ at ~80 ns and then again decline to 118 nm^2^ at the end of simulation. In case of hNatD-H4_S1C, the value fluctuates between around 115 nm^2^ and 124 nm^2^ for the initial 70 ns time and, then it decreases to 118 nm^2^ at the end of simulation. For the initial 70 ns, the overall SASA value of hNatD in hNatD-H4_R3C was remained below to the SASA value of hNatD in hNatD-H4_S1C. However, the SASA value of hNatD in hNatD-H4_R3C was observed slightly higher (~2 nm^2^) as compared to SASA value of hNatD in hNatD-H4_S1C for the last 30 ns. Finally, the SASA value of hNatD in hNatD-H4_G4S was significantly altered (~115 nm^2^-126 nm^2^) over the course of 100 ns simulation which reveals large structural deviation of hNatD as compared to in other complexes. The average Rg values of hNatD were observed 1.74 nm, 1.74 nm, 1.75 nm, 1.75 and 1.76 in hNatD-H4_WT, hNatD-H4_S1C, hNatD-H4_R3C, hNatD-H4_G4D and hNatD-H4_G4S respectively.

To determine the fluctuation of each residue in hNatD, root mean square fluctuation (RMSF) of Cα atoms in residues was calculated for the all complexes. The RMSF value gives details about the interacting residues of hNatD with acetyl-CoA or histone H4 peptide. RMSF is calculated by employing following equation;

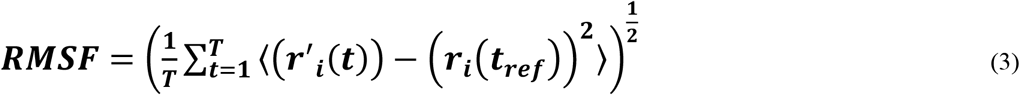

Where, *T* is the simulation time (time steps), *r_i_* is the position of atom *i* at reference time, *t_ref_* and, *r*′ indicates the position of the selected atom at time *t* after aligning with the reference time frame. Fig. 7D shows the RMSF plots of hNatD. Highly dynamic residue regions of hNatD are illustrated in Fig. S3-S7 in hNatD-H4_WT, hNatD-H4_S1C, hNatD-H4_R3C, hNatD-H4_G4D and hNatD-H4_G4S complexes respectively. It can be seen from the RMSF plots that two regions, β6-β7 loop (~180-198 a.a.) at the C-terminal and α1-α2 loop (~68-81 a.a.) showed the highest fluctuation compared to other dynamic regions. hNatD residues in hNatD-H4_G4D have the greater fluctuations whereas residues in hNatD-H4_R3C have the lowest degree of fluctuations. Five dynamic regions, N-terminal α0-helix (~1-20 a.a.), α1-α2 loop (~68-81 a.a.), α2-helix (~81-90 a.a.), α4-helix (~166-176 a.a.) and β6-β7 loop (~180-198 a.a.) in hNatD are part of the catalytic site in which histone H4 tai and acetyl-CoA are located. Protein stability is attributed to the intramolecular hydrogen bonds network and this network provides stability to the protein folds (58). To investigate how WT and MT histone H4 peptides bindings to the hNatD disrupt intramolecular hydrogen bond network in hNatD, we calculated total number of hydrogen bond formation in each complex during the 100 ns time. Fig. 7E shows average number of hydrogen bonds in five complexes. Fig. S8 represents the intramolecular hydrogen bonds variation during 100 ns simulations of studied five complexes. The average number of intramolecular hydrogen bonds were found 162, 162, 167, 158 and 158 in hNatD of hNatD-H4_WT, hNatD-H4_S1C, hNatD-H4_R3C, hNatD-H4_G4D and hNatD-H4_G4S complexes respectively. Thus, hNatD in hNatD-H4_R3C has the greater stability and hNatDs in hNatD-H4_G4D and hNatD-H4_G4S have the least stability. However, hNatDs in hNatD-H4_WT and hNatD-H4_S1C have the stability between prior complexes. Hence, it can be concluded that R3C mutation in histone H4 peptide enhances the protein stability through increasing the number of hydrogen bonds in hNatD whereas, G4D and G4S mutations significantly disturb the hNatD hydrogen bond network and destabilize hNatDs. Also, we obtained intermolecular hydrogen bonds between H4/acetyl-CoA and hNatD during entire simulation in five complexes (Fig. S9). The average number of hydrogen bonds between H4 and hNatD was found 6, 6, 8, 6 and 6 in hNatD-H4_WT, hNatD-H4_S1C, hNatD-H4_R3C, hNatD-H4_G4D and hNatD-H4_G4S complexes respectively (Fig. S9A-E). Thus, R3C mutant H4 peptide has greater stability inside the catalytic site in comparison with other WT and MT histones. And, the average number of hydrogen bonds between acetyl-CoA and hNatD was observed 9, 7, 8, 7 and 6 in hNatD-H4_WT, hNatD-H4_S1C, hNatD-H4_R3C, hNatD-H4_G4D and hNatD-H4_G4S complexes respectively (Fig. S9F-J). Hence, the stability of acetyl-CoA in WT and S1C complexes is greater as compared to other complexes. Thus, this can be considered governing cause for the reduction in catalytic efficiency of hNatD in mutant complexes.

Furthermore, we generated normalized linear mutual information (nLMI) correlation metrics to probe the inter-residues communication network in hNatD under the influence of histone H4 peptide binding. The following equation is used to calculate the LMI between the residue i and j;

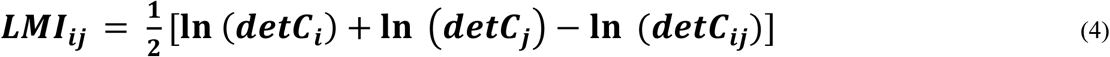

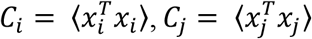, and *C_ij_* = 〈(*x_i_*, *x_j_*)^*T*^〉 where *x_i_* = *R_i_* – 〈*R_i_*〉 and *X_j_* = *R_j_* – 〈R_j_)〉 in which *R_i_*, and *R_j_* are the position vectors of atom *i* and *j*. For the calculation, residues which have LMI ≥ 0.3 and distance threshold ≤ 0.7 Å were considered to be interacting with each other in the protein. In normalized LMI correlation, it gives 0 value for LMI in case of no correlation and 1 when atoms are completely correlated.

To understand the impacts of WT and MTs H4 peptides on hNatD residue network during the simulation, we performed linear mutual correlation analysis on the MD trajectories. The nLMI plots are given in Fig. S10. It is apparently clear from Fig. S10 that among five complexes of histone H4 peptide (WT & MTs) with hNatD, there is considerably lower correlation between residues in hNatD of hNatD-H4_R3C complex as compared to other complexes. As depicted in Fig. S10 that five significantly fluctuating parts (α4-helix, β6-β7 loop, α1-α2 loop, α2-helix and N-terminal α0-helix) of hNatD in five complexes have remarkable differences in correlation.

Additionally, we calculated the correlation difference in hNatD of WT and MT complexes (Fig. 8). It can be clearly seen from the Fig. 8 that MTs have significant correlation difference. The large correlation difference was observed in hNatD-H4_R3C. In this complex, a large number of residues show slightly positive correlation and some residues are appeared to be having higher correlation (Fig. 8B). Thus, residues communicate cooperatively in this complex. However, in other MTs (S1C, G4D and G4S) complexes, anticorrelation motions were observed (Fig. 8A,C-D). This indicates that in these MTs, the residue network of the hNatD is disrupted significantly. Recently, Ho et al. have investigated the catalytic efficiency of hNatD for pentapeptide of H4 WT and MT peptides (17). Their finding shows that the catalytic efficiency of N-α-acetylation of MT histone tails was remarkably reduced. The *K_m_* values for WT, S1C, R3C, G4D and G4S were found 5.9 μM, ~500 μM, 17 μM, 272 μM and 14 μM respectively. The higher values of *K_m_* for S1C and G4D may be due to the significant disruption of hNatD structure as it has anticorrelation residue networks in these two complexes. And, the lower values of *K_m_* for another two mutants, R3C and G4S may be considered due to absence of anticorrelation motions. The WT, R3C and G4S histone tails do not alter the dynamic residue network of catalytic site markedly thus, their catalytic efficiencies were preserved largely as compared to S1C and G4D tails. Hence, our molecular level investigation supports the previously reported finding.

**FIGURE 8.**
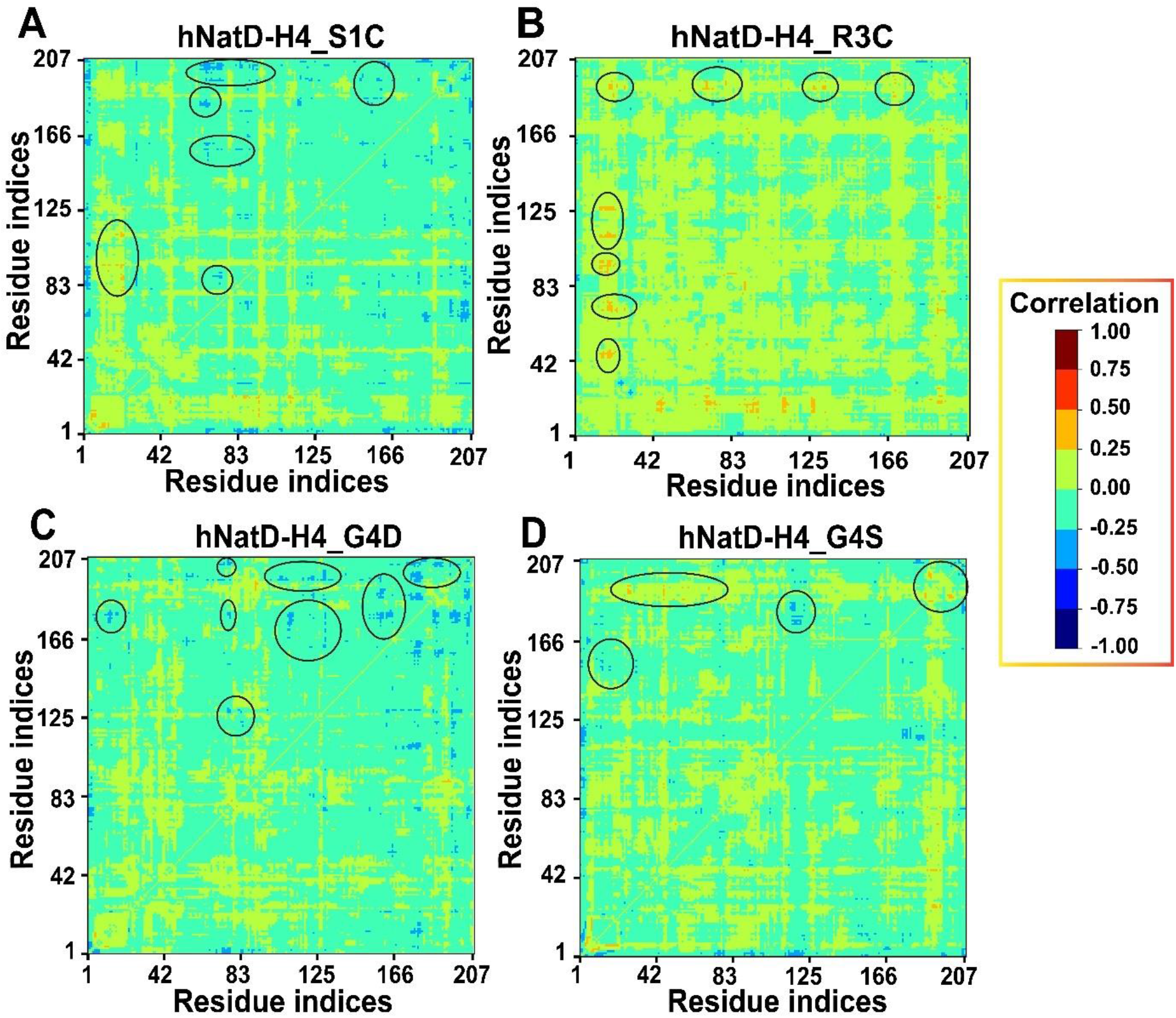
The nLMI correlation difference between WT and MTs complexes of hNatD and histone H4. (A) hNatD-H4_S1C (B) hNatD-H4_R3C (C) hNatD-H4_G4D (D) hNatD-H4_G4S. The degree of correlation corresponds to the color bar.

Furthermore, we calculated betweenness centrality (BC) in complexes. The centrality analysis indicates the degree of communication between residues. The higher BC values reveal protein region where significant inter-domain communication would be present (59). From Table 1, considering all residues which are in close proximity inside the catalytic site, we determined position of every residue in various regions of hNatD in five complexes (Table S2). It was noted that regions which are part of catalytic site have substantial communication (Fig. 9). Most notable regions such as β3, β4, β5 and β7 have been found to have significant connection in every complex whereas only some number of residues from the α0, α1, α2 and α3 are communicate with former regions. Apart from these, residues from the loops (α1 -α2, α2-β2, α3-β5, α4-β5 and β6-β7) have notable communication with former regions. Also, there are only six residues (R112, F113, Q128, L160, I200 and S202) common among five complexes and other are different in each complex thus, mutations lead remarkable changes in the hNatD structure.

**TABLE 1.**
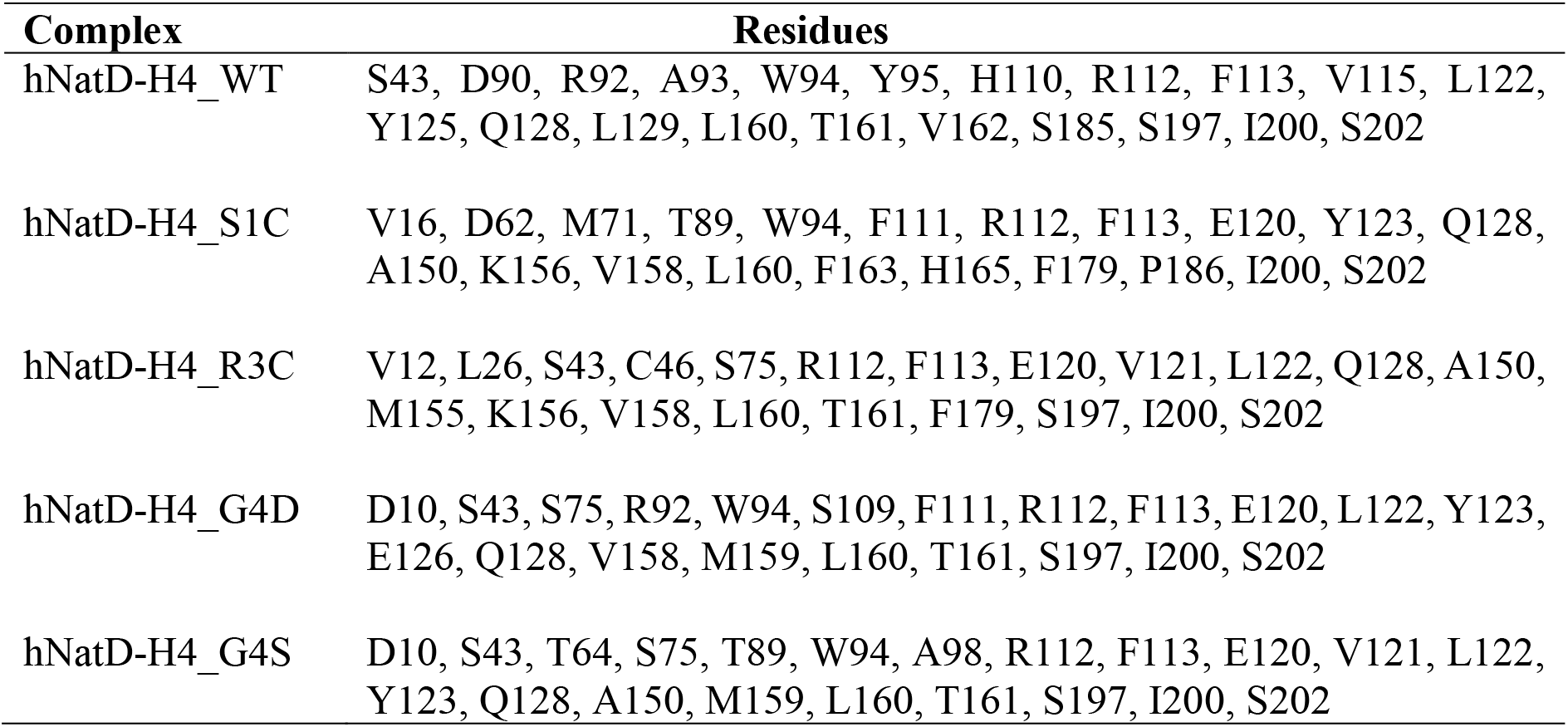
The large betweenness centrality (BC) containing residues of hNatD in studied five complexes with histone H4 peptides (WT & MTs).

**FIGURE 9.**
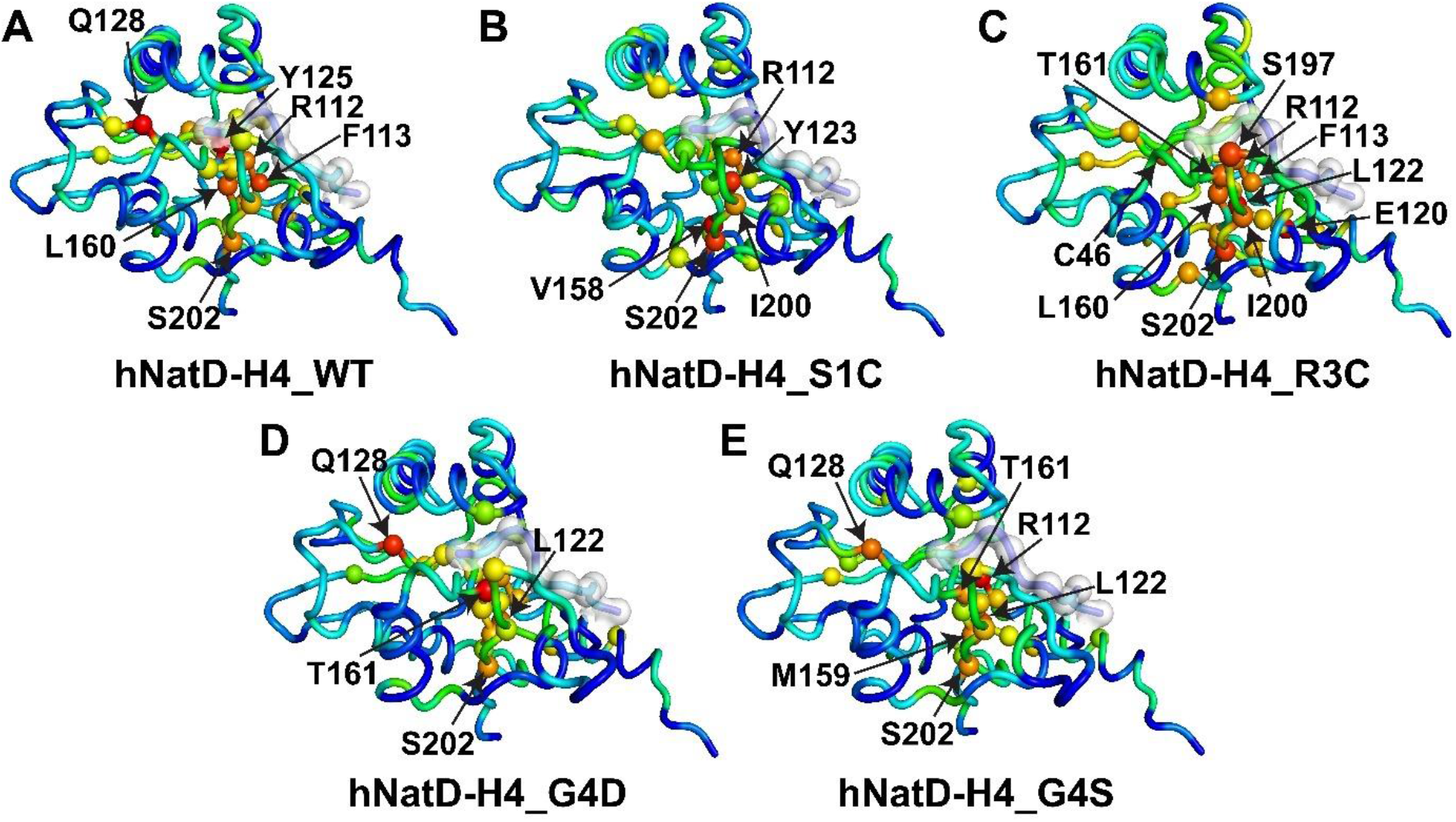
Betweenness centrality (BC) of hNatD in WT and MT complexes. (A) hNatD-H4_S1C (B) hNatD-H4_R3C (C) hNatD-H4_G4D (D) hNatD-H4_G4S. Spheres are the residues which have greater communication in hNatD. Red color has the highest degree of betweenness and blue has the least. Histone H4 peptides are shown in cartoon-surface representation.

Furthermore, in addition to the beta strands and helices, loop regions are also communicating in hNatD. It was noted that there are a greater number of residues show communication from the loop regions in MTs complexes as compared to WT complex of hNatD. The β6-β7 loop communication was observed in all the complexes. In WT, another loop (α2-β2) is part of the communication. However, another type of loops such as α0-β0, α1-α2, α3-β5 and α4-β5 were observed to participate to the communication in MTs complexes. Among these five loops in WT and MTs, except α3-β5 loop every loop is part of acetyl-CoA and substrate recognition sites. Thus, it is clear that during the substrate recognition and acetylation, these beta strands, helices and loops dynamically communicate and work cooperatively. It can be seen from the Fig. 9 that in hNatD-H4_WT and hNatD-H4_R3C, the residue contact network is maintained at the catalytic sites as compared to other MT complexes. Hence, it implies that catalytical core of the hNatD in these two complexes are not influenced greatly.

### INTERACTION NETWORK BETWEEN H4 PEPTIDE AND hNatD

To understand the interaction dynamics between the histone H4 peptides (WT & MTs), we extracted the 10 frames from 10 ns to 100 ns at every 10 ns from the each complex and calculated the interactions and plotted dominant interactions. From the Fig. 2B, we know that the catalytic site of hNatD is highly acidic in nature compared to previous hNatA (13). Thus, dominant interactions come from acidic residues in all the complexes. Results indicate that as compared to hNatD-H4_WT, the total number of stable interactions in MTs were observed large in number. And, this is the cause of significant deflection of WT histone tail inside the catalytical core in hNatD (Fig. 5A). In MTs, greater number of residues interact with the histone H4 peptide and stabilize it inside the hNatD (Fig. 5B-E).

In hNatD-H4_WT, histone H4 peptide interacts with E83 and E87 from α2-helix, D90 from α2-β2 loop and, D114, V115 and E116 from the β3 strand through S1, R3 and K5 residues (Fig. 10A). The hydrogen bonds and salt bridges are the dominant types of interactions between H4 peptide and hNatD. E83 forms hydrogen bonds and salt bridges with S1 hydroxyl and R3 amino groups. Also, E87 interacts to S1 hydroxyl and R3 amino groups through forming hydrogen bonds and salt bridges with amino groups of S1 and R3 residues. The K5 sidechain amino group interacts with the D115, V115 and E116 hNatD residues by forming hydrogen bonds and salt bridges.

**FIGURE 10.**
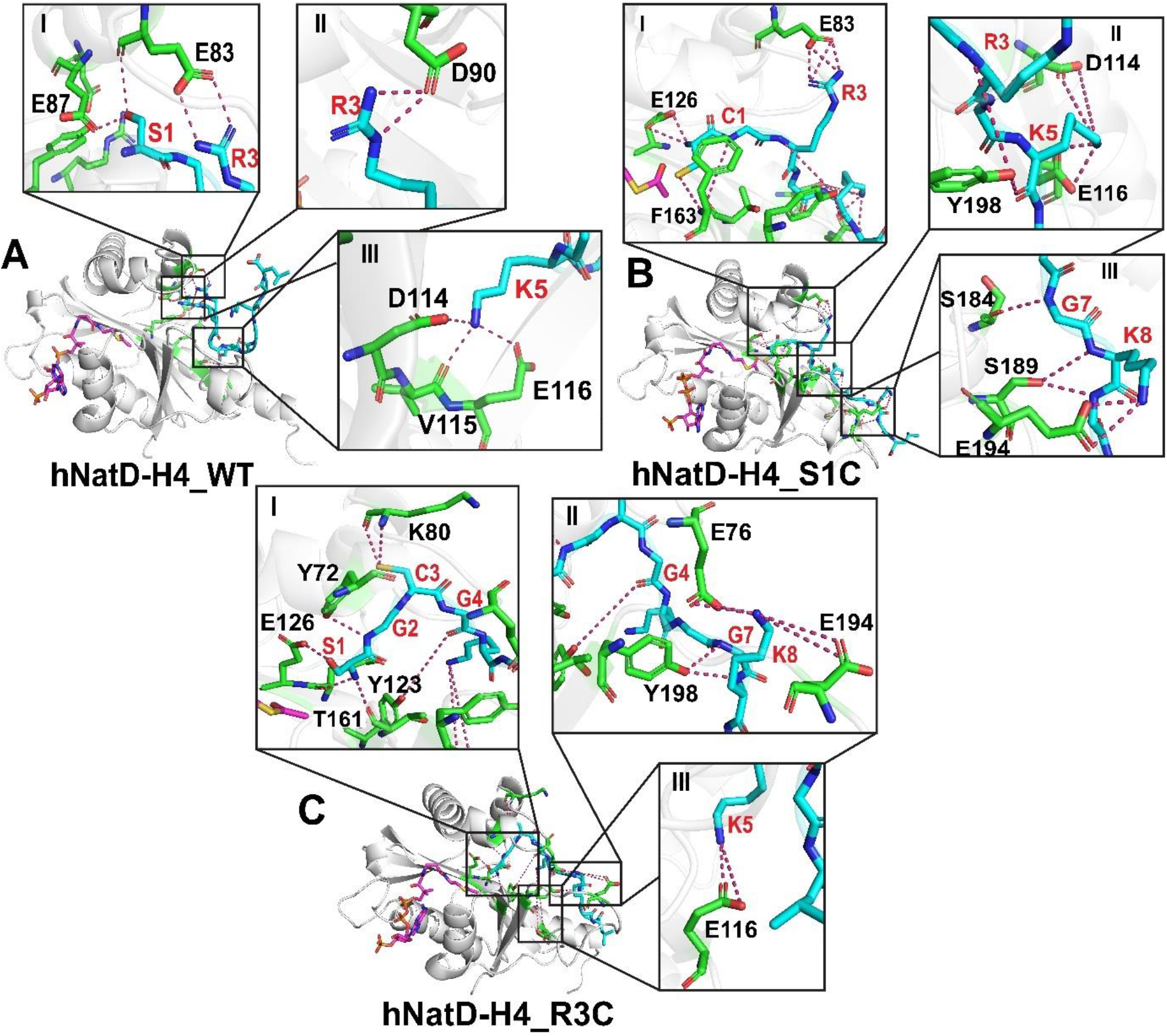
Interactions between histone H4 peptides and hNatD. (A) hNatD-H4_WT (B) hNatD-H4_S1C (C) hNatD-H4_R3C. Histone H4 residues are colored in red while hNatD residues are in black.

In MTs, new residues from the hNatD appear and increase the binding affinity of H4 peptide with hNatD. Thus, the interaction network is largely altered in MTs in comparison with WT. The most important residue, E126 was observed to be participating in interactions with all MT H4 peptides and this might be cause of the stability of MT H4 peptides at the binding site. In case of hNatD-H4_S1C, E116 became dominant and E194 from the β6-β7 loop was emerged to interact with H4 peptide. In addition to that, two aromatic residues (F163 of β5 strand and Y198 of β7 strand) were also found to have interactions with H4 peptide (Fig. 10B). In this case, E83 only interacts to R3 by forming hydrogen bonds and salt bridges. And, E126 of β4 strand forms hydrogen bonds and salt bridges with the amino groups of C1 residue. The thiol group in C1 also forms hydrogen bonds with F163 residue. The V115 residue was disappeared and new residue, Y198 was observed to interact with the R3 and K5 through forming hydrogen bonds. Like E116, D114 also interacts with the K5 sidechain through hydrogen bonds and salt bridge. Furthermore, S184 and S189 from the β6-β7 loop bind to the G7 and K8 of the H4 peptide by forming hydrogen bonds.

In hNatD-H4_R3C, E83 was vanished and new acidic residue E76 from the α1-α2 loop was found to involved in interactions (Fig. 10C). E76 interacts to K8 residue of the H4 peptide by forming hydrogen bond and salt bridge. In this mutation, basic residue, arginine is substituted with cysteine polar residue. Hence, K80 from α1-α2 loop was found to interact with the cysteine thiol group by forming hydrogen bonds. Along with E126, another residue (T161 from β5 strand) was observed to interact with S1 residue of the H4 peptide. Y72 residue from the α1-helix was found to have interaction with G2 of histone tail. Y123 from the β4 strand another new residue that was observed to bind with G4 residue. E116 and E194 interact with K5 and K8 residues of the histone H4 respectively like in hNatD-H4_S1C complex. However, Y198 interacts with G7 and K8 only unlike in hNatD-H4_S1C complex where it interacts with R3 and K5.

At the fourth position in histone H4, there were two mutations studied (G4D and G4S). Fig. 11 illustrates the interactions of G4S and G4D H4 with hNatD. It can be seen from the Fig. 11A-B that there are large number of hNatD residues interact with the G4S H4 as compared to G4D H4 peptide. The G4D mutation causes significant alternation in the hNatD structure due to the acidic sidechain of the aspartic acid. The H4 binding site is highly acidic in nature (Fig. 3B) thus, this carboxylic group repels surrounding residue through electrostatic interactions. Along with hNatD structural alternation, this residue also significantly impacts the position of acetyl-CoA inside the hNatD in comparison with other complexes (Fig. 5D).

**FIGURE 11.**
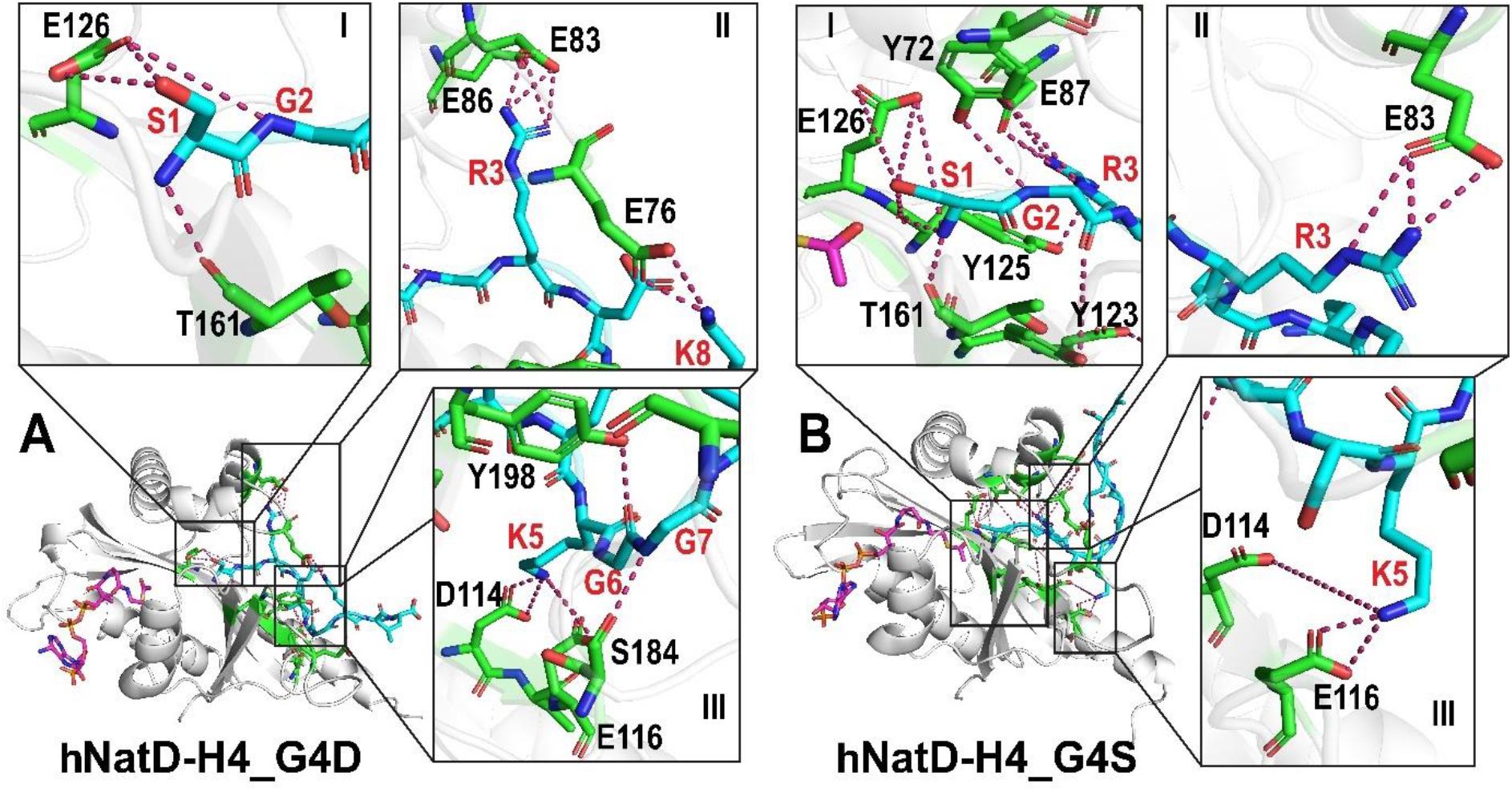
Interactions between histone H4 peptides and hNatD. (A) hNatD-H4_G4D (B) hNatD-H4_G4S. Histone H4 residues are colored in red while hNatD residues are in black.

The commonly interacting residues are E83, D114, E116, E126 and T161. However, E83, D114 and E116 are the dominant residues in hNatD-H4_G4D and, E116 and E126 are in hNatD-H4_G4S. These residues form hydrogen bonds and salt bridges with the MT H4 peptides. In case of hNatD-H4_G4D, E126 interacts with the S1 and G2 through hydrogen bonds only. T161 forms hydrogen bond with the S1 residue. Two acidic residues (E83 and E86) interact to the sidechain of the R3 whereas another acidic residue, E76 interacts with the K8 through hydrogen bond and salt bridge. The K5 residue surrounds with two acidic residues, D114 and E116 and interacts by forming hydrogen bonds and salt bridges. S184 and Y198 interact with the G7 and G6 through only hydrogen bonds. However, there were two aromatic residues, Y72 and Y123 observed to have interactions with G2 and, Y125 with S1 and R3 by forming hydrogen bonds only in hNatD-H4_G4S complex. Like in hNatD-H4_G4D, D114 and E116 interact with the K5 residue of the H4 in hNatD-H4_G4S. The H-bond and salt bridge interactions between H4 peptides and hNatD in five complexes are given in Table S3. Table 2 shows the interacting residues of hNatD with WT and MT H4.

**TABLE 2.**
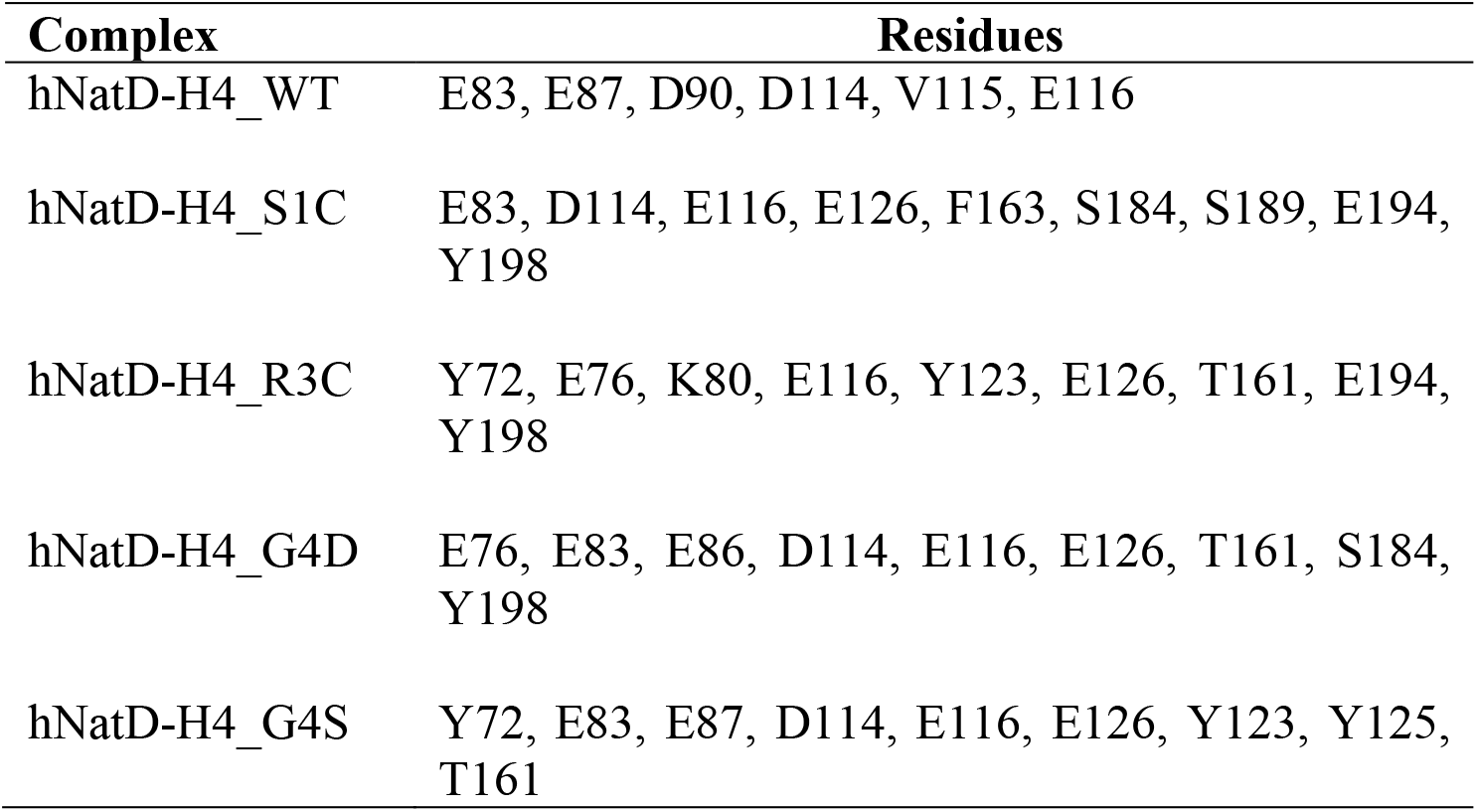
Interacting residues of hNatD with H4 (WT & MT) in five complexes.

Also, we calculated hydrogen bond, electrostatic and van der Waals energies of the WT and MTs complexes to investigate the binding energetics. These three energies were calculated for every 10 ns frame from the 10 ns to 100 ns in each complex (Fig. 12). The total stabilizing energy of the complex was calculated using adding all three energies. These energies are plotted in the form of heatmaps and represented in Fig. 12. The values of these energies are given in Table S4. It can be seen from the Fig. 10 that hNatD-H4_R3C has the highest amount of stabilization energy. The maximum contribution to total stabilization energy comes from the van der Waals and less from the hydrogen bond energy. The hydrogen bond, and van der Waals energies in MT complexes are large as compared to WT complex. However, electrostatic energy was observed higher in WT complex.

**FIGURE 12.**
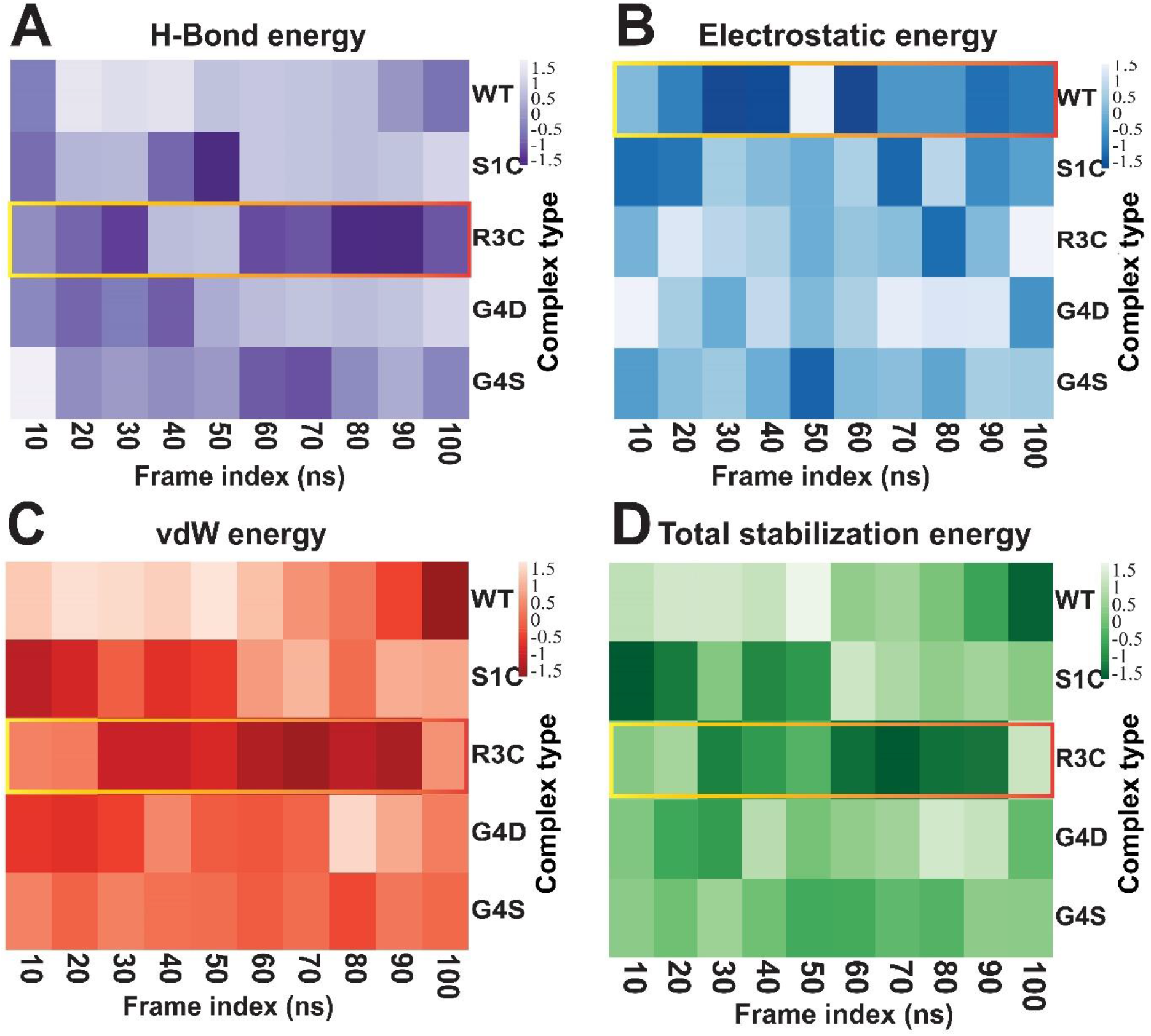
Heatmaps of energy contributions (kJ/mol) to the interactions calculated using PIMA web server. (A) Hydrogen bond energy (B) Electrostatic energy (C) van der Waals energy (D) Total stabilization energy. The yellow-red rectangular box shows the overall greater energy contribution to the complex.

## CONCLUSION

In this study, we examined the impacts of WT and MTs (S1C, R3C, G4D and G4S) histone H4 decapeptides on their bindings with hNatD by using 100 ns all-atom MD simulation. Our results support previous finding that the mutant H4 histones reduce the catalytic efficiency of hNatD. The MD post-trajectory analyses revealed that S1C, G4S and G4D mutants remarkably alter the residue network in hNatD. The intramolecular hydrogen bond analysis suggested that there is considerable number of losses of hydrogen bonds in hNatD of hNatD-H4_G4D and hNatD-H4_G4S complexes whereas large number of hydrogen bonds were increased in hNatD of hNatD-H4_R3C complex during the entire simulations. This implies that R3C mutant binding to hNatD brings stability in hNatD in comparison with WT and other MTs complexes. The linear mutual information (LMI) and Betweenness centrality (BC) suggest that S1C, G4D and G4S significantly disrupt the catalytic site residue network as compared to R3C mutation in H4 histone. Thus, this might be the cause of notable reduction in catalytic efficiency of hNatD in these three mutant complexes. Further, interaction analysis supports that E126 is the important residue for the acetyltransferase mechanisms as it is dominantly found to have interactions with numerous residues of MTs histones in MD frames. Additionally, intermolecular hydrogen bond and RMSD analyses of acetyl-CoA predict the higher stability of acetyl-CoA inside the WT complex of hNatD and R3C complex. Also, we report here structural and dynamic aspects and residue interactions network (RIN) of hNatD to target it to control the cell proliferation in the lung cancer conditions.

## ACKNOWLEDGEMENTS

Shravan B. Rathod is thankful to his Chemistry Department for providing computational and infrastructure facilities. He is also thankful to Dr. Mustafa Tekpinar from the Pasteur Institute, France for explaining correlation plots and valuable suggestions on draft.

## AUTHOR CONTRIBUTIONS

**Shravan B. Rathod:** Conceptualization, Investigation, Methodology, Data curation, Formal analysis, Writing-original draft, Writing-review & editing. **Kinshuk Raj Srivastava:** Conceptualization, Investigation, Supervision, Methodology, Data curation, Formal analysis, Writing-original draft, Writing-review & editing.

