## Supplemental File for "Dynamic play between human N-α-acetyltransferase D and H4-mutant histones: Molecular dynamics study" (from another version of this preprint: version 1, 2022-03-18)

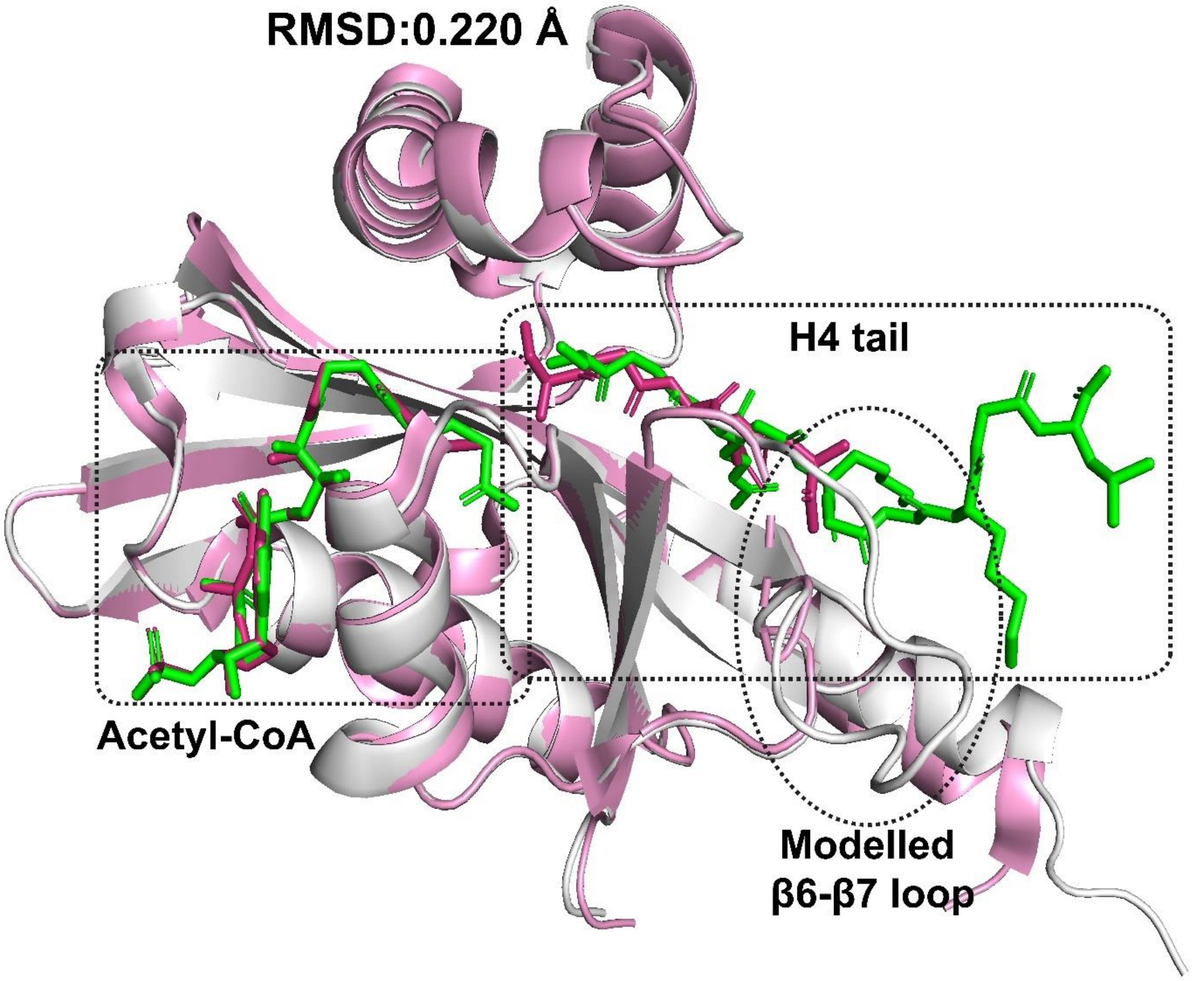


**Figure S1.** Structural alignment of crystal structure of hNatD (PDB: 4U9W) with docked structure. Crystal and docked structures are colored in light pink and white respectively. Modelled Acetyl-CoA and histone H4 tail are shown in green color whereas crystal structures are represented in pink color.

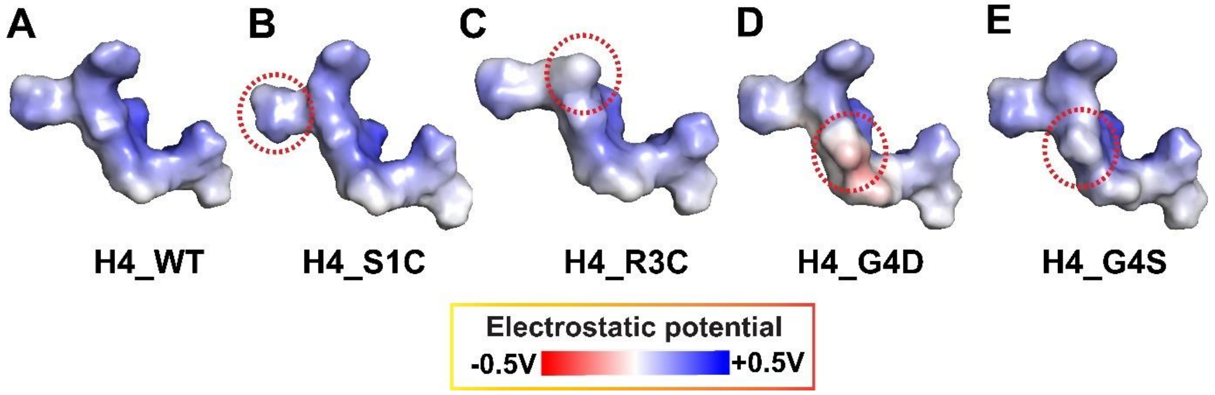


**Figure S2.** Electrostatic potential surface representations of WT and MTs histone H4. (A) H4_WT (B) H4_S1C (C) H4_R3C (D) H4_G4D (E) H4_G4S. Red colored dashed circle indicates the position of mutation.

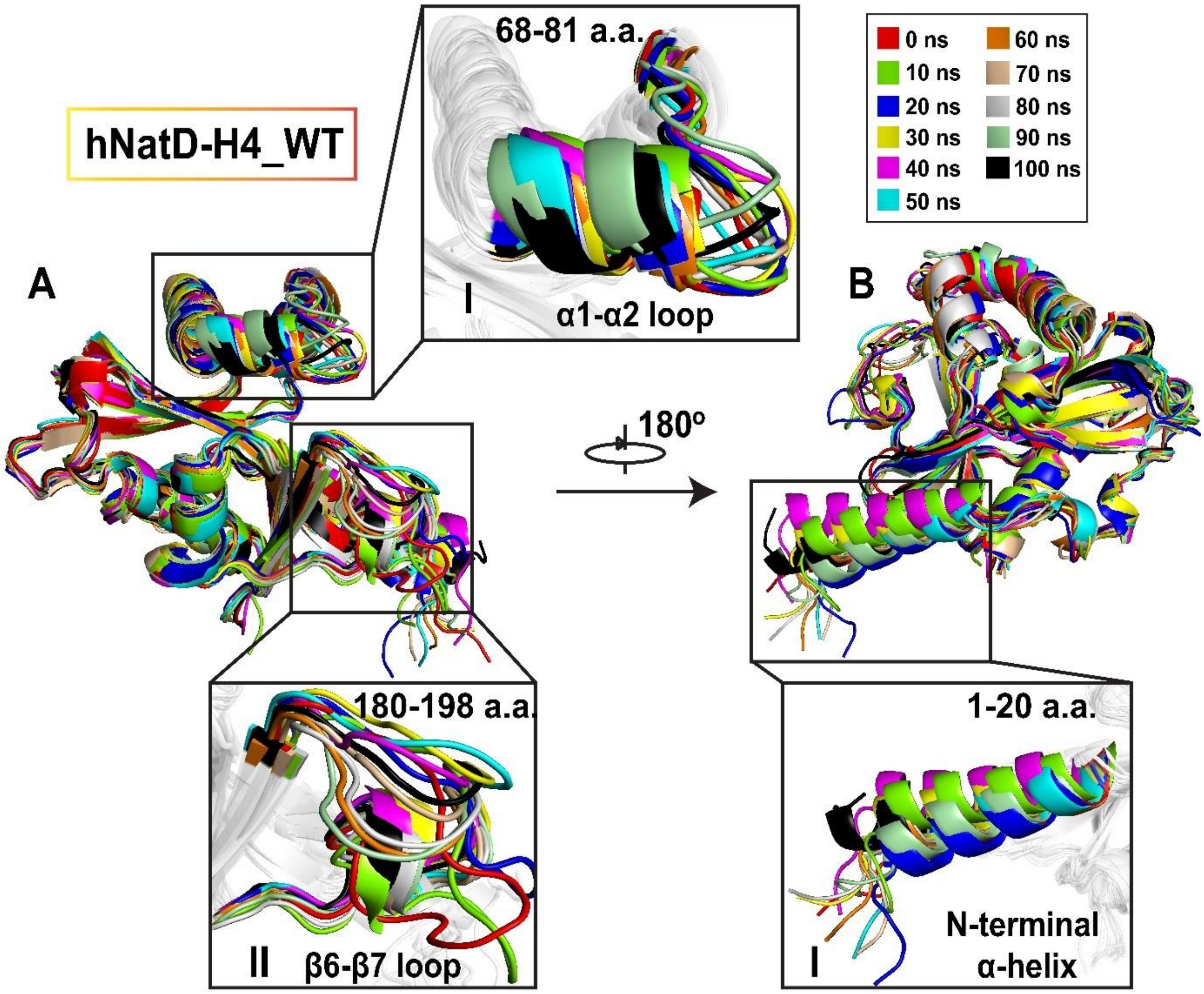


**Figure S3.** Structural alignment of MD trajectory frames of hNatD from NatD-H4_WT complex at every 10 ns from 0 ns to 100 ns. Highly dynamics parts of the hNatD are shown in square boxes.

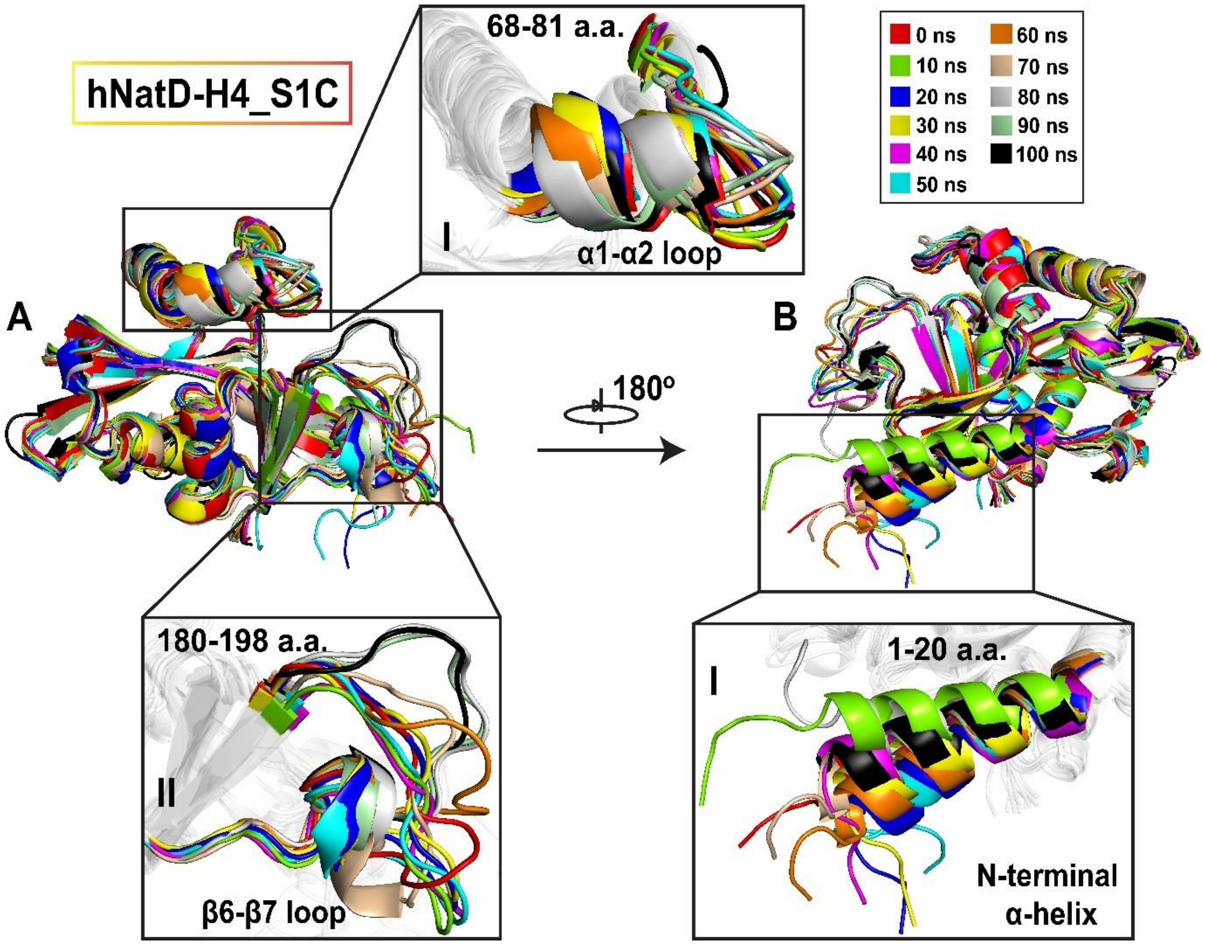


**Figure S4.** Structural alignment of MD trajectory frames of hNatD from hNatD-H4_S1C complex at every 10 ns from 0 ns to 100 ns. Highly dynamics parts of the hNatD are shown in square boxes.

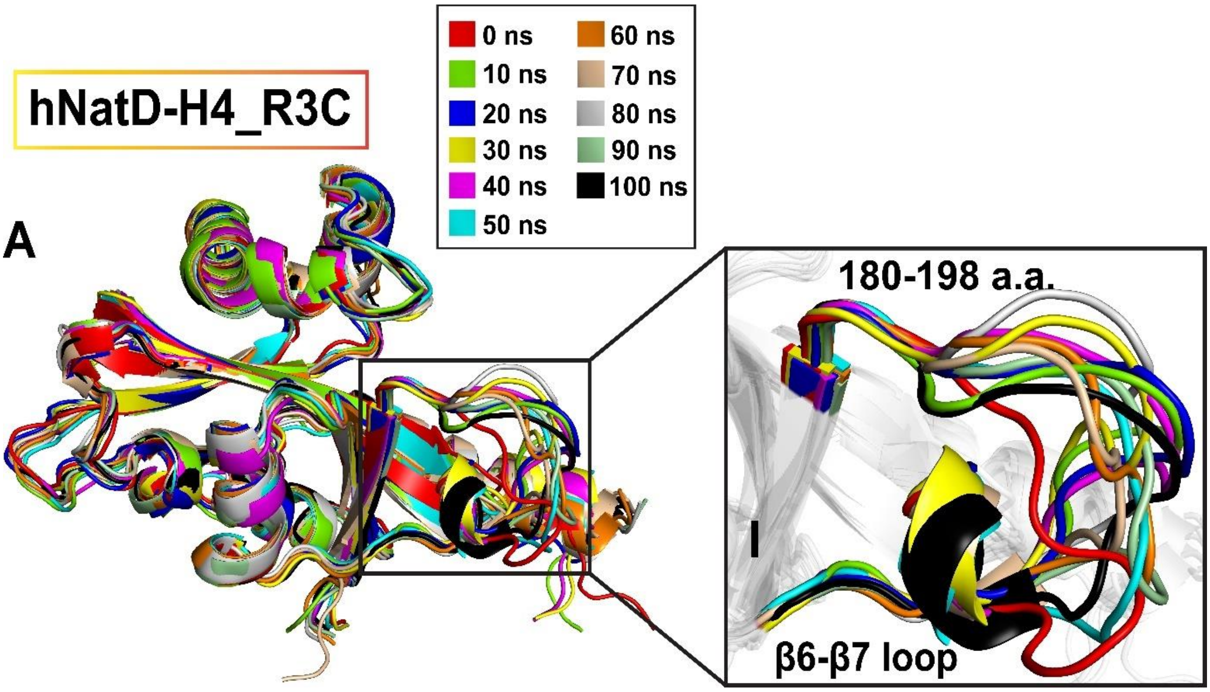


**Figure S5.** Structural alignment of MD trajectory frames of hNatD from hNatD-H4_R3C complex at every 10 ns from 0 ns to 100 ns. Highly dynamics parts of the hNatD are shown in square boxes.

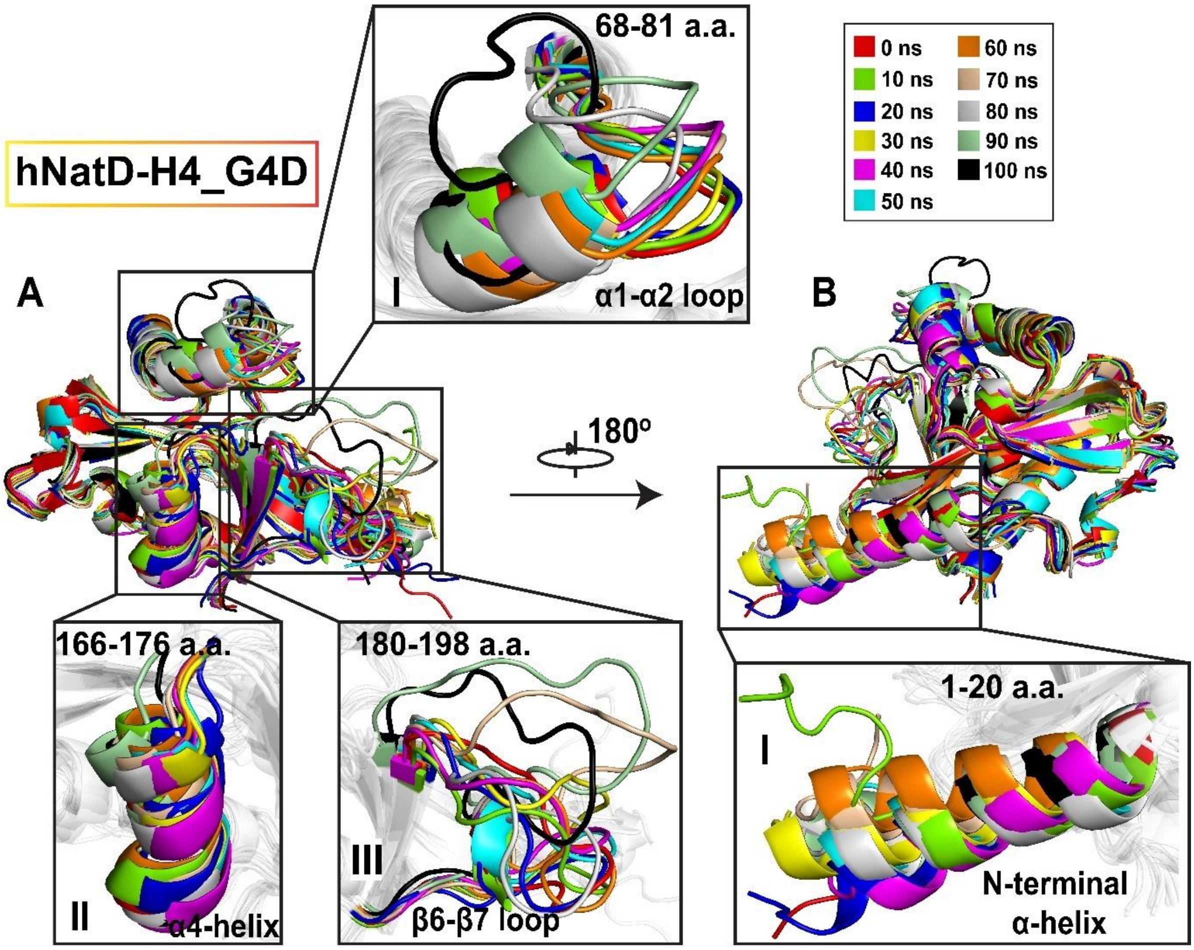


**Figure S6.** Structural alignment of MD trajectory frames of hNatD from hNatD-H4_G4D complex at every 10 ns from 0 ns to 100 ns. Highly dynamics parts of the hNatD are shown in square boxes.

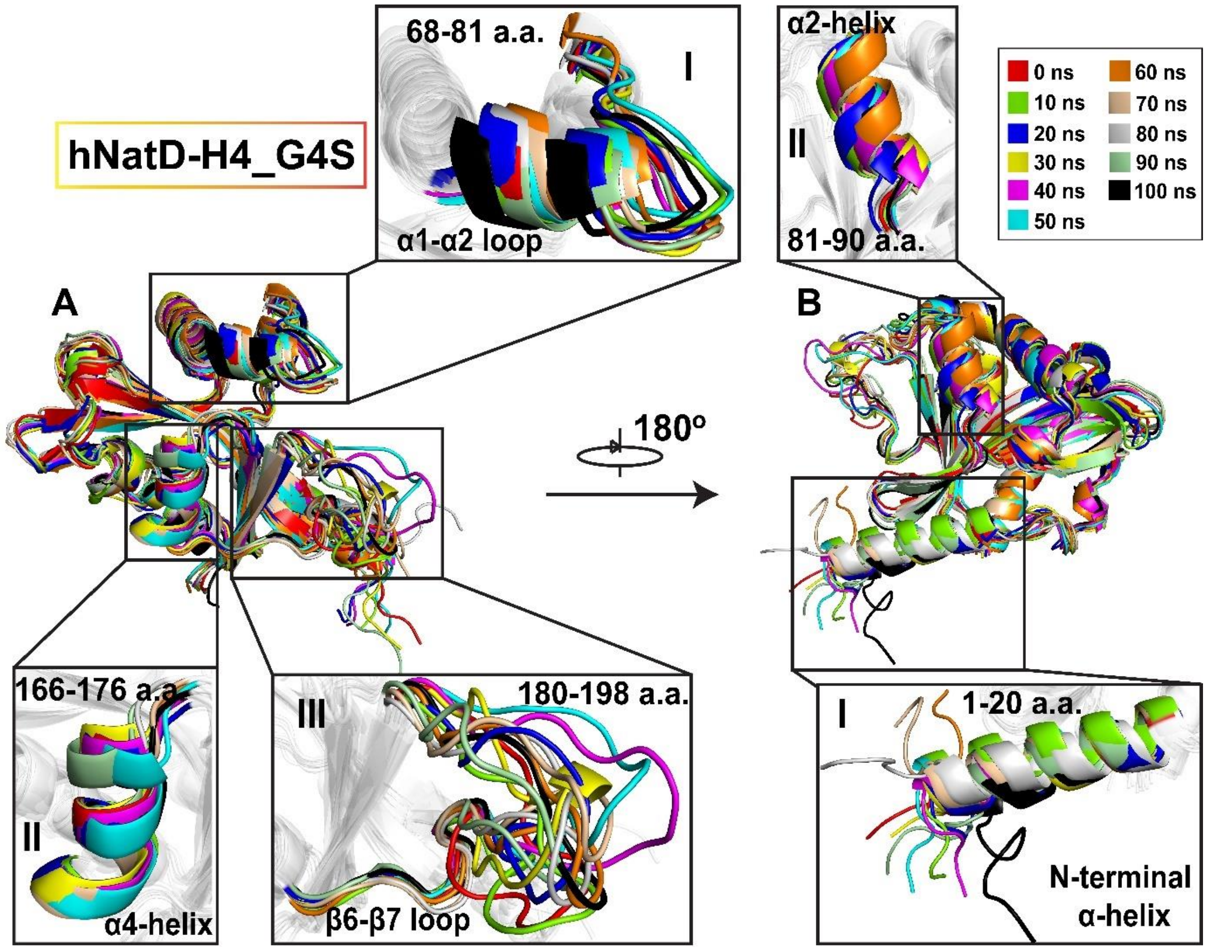


**Figure S7.** Structural alignment of MD trajectory frames of hNatD from hNatD-H4_G4S complex at every 10 ns from 0 ns to 100 ns. Highly dynamics parts of the hNatD are shown in square boxes.

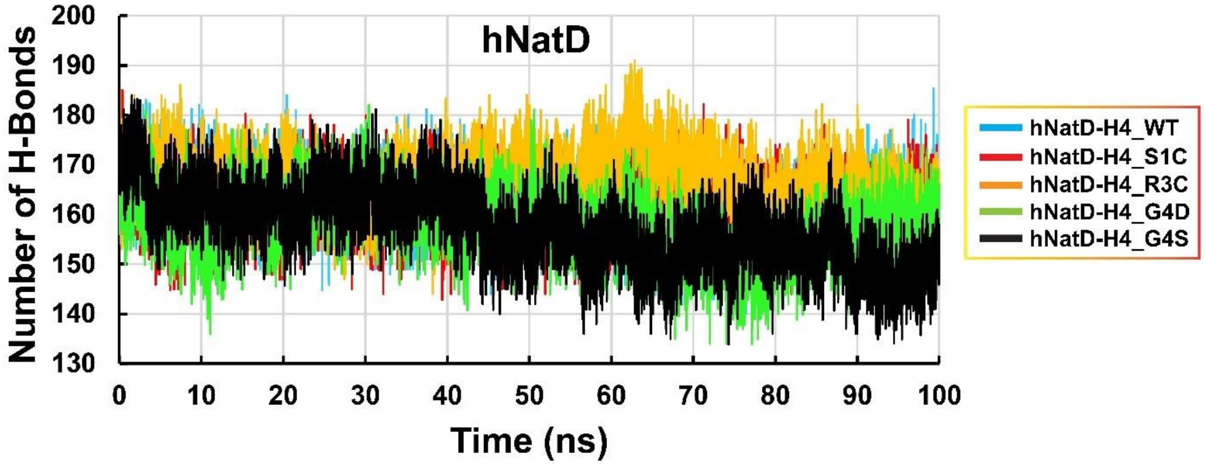


**Figure S8.** Intramolecular hydrogen bonds in hNatD during 100 ns simulations of five complexes.

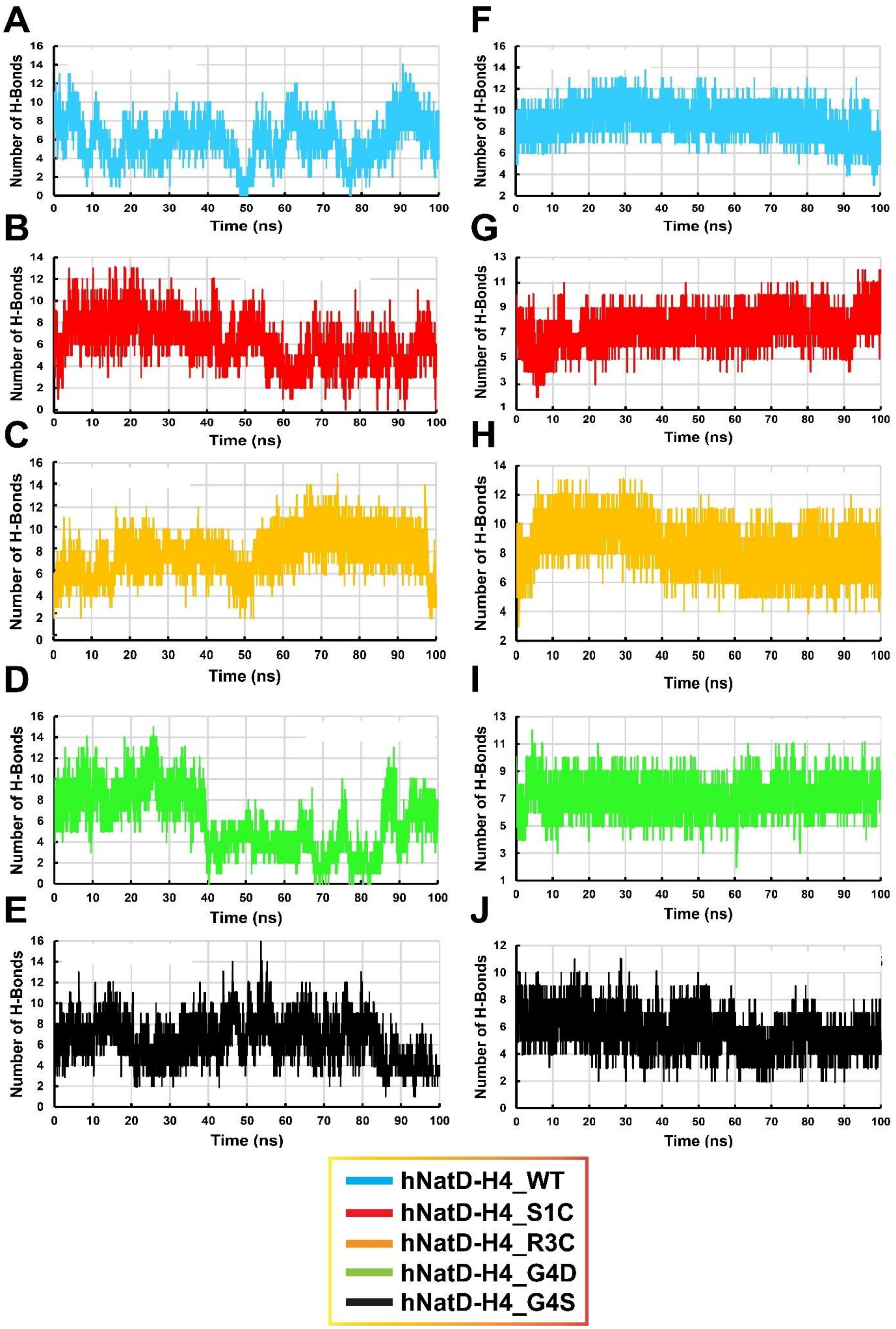


**Figure S9.** Intermolecular hydrogen bonds in five complexes during 100 ns simulations. (A-E) Hydrogen bonds between H4 histone and hNatD. (F-J) Hydrogen bonds between acetyl-CoA and hNatD.

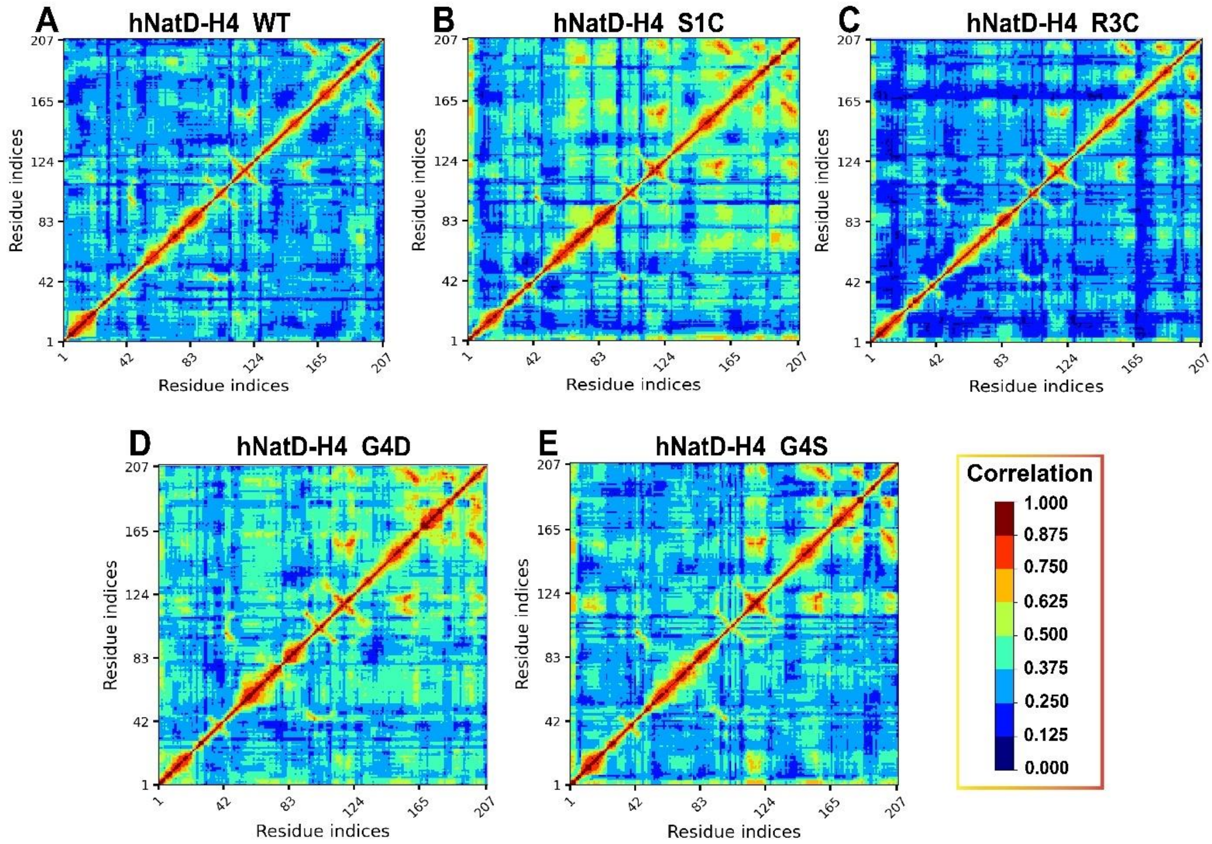


**Figure S10.** nLMI plots of hNatD in WT and MTs complexes of histone H4 tail. (A) hNatD-H4_WT (B) hNatD-H4_S1C (C) hNatD-H4_R3C (D) hNatD-H4_G4D (E) hNatD-H4_G4S. The degree of correlation corresponds to the color bar.

**Table S1.** The *correlationplus* installation and usage commands.

| Task | Commands |
|---|---|
| ***correlationplus* Installation & usage** | *correlationplus* installation guide and video tutorials for use are given at https://github.com/tekpinar/correlationplus |
| ***correlationplus* activation** | #Go to source folder and type following command;<br><br>source bin/activate |
| **nLMI calculation** | correlationplus calculate -p hNatD_WT.pdb -f hNatD_WT.xtc -t nlmi #Use recentred XTC/TRR trajectory<br><br>correlationplus visualize -p hNatD_WT.pdb -i nLMI.dat -t nlmi -o nlmi-correlations |
| **nLMI difference calculation** | correlationplus diffMap -i hNatD_WT-nLMI.dat -j hNatD_MT-nLMI.dat -p hNatD_WT.pdb |
| **Residue betweenness calculation** | correlationplus analyze -t nLMI.dat -p hNatD_WT.pdb -c betweenness<br><br>vmd -e correlation_betweenness.tcl or open correlation_betweenness.pml file in PyMOL. |

**Table S2.** Location of higher Betweenness centrality (BC) containing hNatD residues in five complexes. hNatD regions are color coded according to illustrated in Figure 2. Helices, β-strands and loops are colored in pink, blue and grey colors respectively.

| Complex | Correlated residue position | |
|---|---|---|
| **hNatD-H4_WT** | α2: D90<br>β1: S43<br>β2: W94, Y95<br>β3: H110, R112, F113, V115<br>β4: L122, Y125, Q128, L129 | β5: L160, T161, V162<br>β7: I200, S202<br>α2-β2: D93<br>β6-β7: S185, S197 |
| **hNatD-H4_S1C** | α0: V16<br>α1: D62, M71<br>α2: T89<br>α3: A150<br>β2: W94<br>β3: F111, R112, F113 | β4: E120, Y123, Q128<br>β5: V158, L160, F163<br>β7: I200, S202<br>α3-β5: K156<br>α4-β5: H165<br>β6-β7: F179, P186 |
| **hNatD-H4_R3C** | α0: V12<br>α3: A150<br>β1: S43, C46<br>β3: R112, F113<br>β4: E120, V121, L122, Q128<br>β5: V158, M159, L160, T161 | β7: I200, S202<br>α0-β0: L26<br>α1-α2: S75<br>α3-β5: M155, K156<br>β6-β7: F179, S197 |
| **hNatD-H4_G4D** | α0: D10<br>β1: S43<br>β2: W94<br>β3: S109, F111, R112, F113<br>β4: E120, L122, Y123, E126, Q128 | β5: V158, M159, L160, T161<br>β7: I200, S202<br>α1-α2: S75<br>α2-β2: R92<br>β6-β7: S197 |
| **hNatD-H4_G4S** | α0: D10<br>α1: T64<br>α2: T89<br>α3: A150<br>β1: S43<br>β2: W94, A98 | β3: R112, F113<br>β4: E120, V121, Y123, Q128<br>β5: M159, L160<br>β7: I200, S202<br>α1-α2: S75<br>β6-β7: S197 |

**Table S3.** Calculated H-Bond and salt bridge interactions in every 10 ns frame from 10 ns to 100 ns simulations of five complexes. Also, residue and its overall occurrence was calculated from the total number of interactions. Check Table S3 excel document for this data.

**Table S4.** Calculated four types of energies (H-Bond, electrostatic, vdW and total stabilization) in every 10 ns frame from 10 ns to 100 ns simulations of five studied complexes.

| | H-Bond energy (kJ/mol) | | | | | Electrostatic energy (kJ/mol) | | | | |
|---|---|---|---|---|---|---|---|---|---|---|
| **Frame indices** | **hNatD-H4_WT** | **hNatD-H4_S1C** | **hNatD-H4_R3C** | **hNatD-H4_G4D** | **hNatD-H4_G4S** | **hNatD-H4_WT** | **hNatD-H4_S1C** | **hNatD-H4_R3C** | **hNatD-H4_G4D** | **hNatD-H4_G4S** |
| 10 | -21.85 | -23.06 | -20.36 | -20.83 | -11.01 | -38.1 | -63.3 | -40.4 | -11.9 | -48.3 |
| 20 | 0.00 | -8.64 | -20.00 | -20.11 | -14.43 | -78.8 | -82.9 | -29.0 | -45.4 | -53.3 |
| 30 | 0.00 | -9.41 | -31.98 | -22.61 | -15.38 | -86.0 | -39.9 | -33.8 | -53.3 | -42.0 |
| 40 | 0.00 | -21.16 | -7.38 | -22.56 | -15.38 | -75.6 | -36.4 | -26.1 | -20.5 | -42.0 |
| 50 | 0.00 | -33.01 | 0.00 | -5.17 | -9.70 | -7.2 | -38.7 | -38.7 | -34.6 | -64.1 |
| 60 | 0.00 | 0.00 | -33.00 | -3.68 | -29.65 | -88.0 | -38.4 | -43.7 | -37.4 | -49.4 |
| 70 | 0.00 | 0.00 | -23.40 | 0.00 | -24.17 | -55.9 | -67.4 | -45.5 | -28.2 | -44.9 |
| 80 | 0.00 | 0.00 | -31.80 | 0.00 | -12.09 | -38.3 | -11.6 | -53.8 | -2.1 | -33.2 |
| 90 | -14.12 | 0.00 | -43.03 | 0.00 | -7.28 | -65.1 | -56.8 | -41.7 | -21.6 | -33.9 |
| 100 | -12.96 | 0.00 | -15.95 | 0.00 | -10.69 | -79.3 | -58.6 | 0.0 | -66.3 | -32.7 |
| | **vdW (kJ/mol)** | | | | | **Total stabilization energy (kJ/mol)** | | | | |
| **Frame indices** | **hNatD-H4_WT** | **hNatD-H4_S1C** | **hNatD-H4_R3C** | **hNatD-H4_G4D** | **hNatD-H4_G4S** | **hNatD-H4_WT** | **hNatD-H4_S1C** | **hNatD-H4_R3C** | **hNatD-H4_G4D** | **hNatD-H4_G4S** |
| 10 | -81.3 | -159.6 | -110.6 | -141.3 | -110.3 | -141.3 | -246.0 | -171.4 | -174.0 | -169.6 |
| 20 | -43.1 | -156.1 | -103.5 | -148.4 | -118.5 | -121.9 | -247.7 | -152.5 | -213.9 | -186.2 |
| 30 | -58.0 | -148.5 | -200.8 | -169.3 | -124.2 | -144.0 | -197.8 | -266.6 | -245.2 | -181.6 |
| 40 | -72.7 | -157.1 | -172.1 | -110.7 | -124.2 | -148.2 | -214.7 | -205.6 | -153.8 | -181.6 |
| 50 | -58.1 | -119.4 | -127.3 | -108.6 | -102.4 | -65.3 | -191.1 | -165.9 | -148.3 | -176.2 |
| 60 | -65.8 | -81.8 | -150.1 | -113.3 | -110.8 | -153.9 | -120.2 | -226.9 | -154.4 | -189.8 |
| 70 | -72.5 | -52.7 | -164.2 | -100.2 | -95.9 | -128.4 | -120.1 | -233.0 | -128.4 | -165.0 |
| 80 | -108.3 | -114.0 | -164.4 | -56.7 | -131.4 | -146.7 | -125.6 | -250.0 | -58.8 | -176.6 |
| 90 | -108.1 | -44.0 | -151.5 | -45.8 | -78.2 | -187.4 | -100.9 | -236.2 | -67.4 | -119.3 |
| 100 | -117.4 | -77.2 | -81.6 | -86.5 | -91.0 | -209.6 | -135.8 | -97.6 | -152.8 | -134.4 |
